# 75 years of anthropogenic change and its impact on Canadian butterfly taxonomic and phylogenetic diversity

**DOI:** 10.1101/2023.09.30.560336

**Authors:** J.M.M Lewthwaite, A.Ø Mooers

## Abstract

Previous studies have documented very little net change in average quadrat-level species richness and phylogenetic diversity. However, although the average remains centered around 0, there is much variation around this mean and many outliers. The relative contribution of anthropogenic drivers (such as climate change or land use change) to these outliers remains unclear. Traits may dictate species responses to these changes, and if relatedness is correlated with trait similarity, then the impacts of anthropogenic change may be clustered on the phylogeny. We build the first regional phylogeny of all Canadian butterfly species in order to examine change in community phylogenetic structure in response to two main documented drivers of change -- climate change and land use change -- across 265 species, 75 years and 96 well-sampled quadrats. We find no evidence that, on average, communities are becoming more or less clustered than one would expect. However, there is much variation depending on the magnitude and type of anthropogenic change occurring within a quadrat. We find that climate change as well as agricultural development is reducing species richness within a quadrat, and these species that are lost tend to be scattered across the phylogeny. However, agricultural abandonment is having the opposite effect: we find increasing species richness in the years immediately following it and decreasing distance between species in quadrats with the highest rates of abandonment, such that the species that colonize these plots tend to be close relatives of those already present and thus contribute little novel phylogenetic diversity to an assemblage. Consistent with previous work, small changes in local species richness may conceal simultaneous change in other facets of biodiversity.

## Introduction

Several studies have recently documented little to no net change in plot-level richness (Velland et al. 2013; Dornelas et al., 2014; Elahi et al., 2015; Lewthwaite and Mooers 2021; see Velland et al. 2017 for a review) and abundance through time (Crossley et al. 2020). Though the global average is centered on zero (Dornelas et al. 2019), there is often a high amount of variation around this mean (Blowes et al. 2019).

Various theories and drivers have been proposed for these outliers. Land use change has been most often linked to richness declines (Newbold et al. 2015); a global meta-analysis found that land use change has resulted in a 24.8% decline in species richness across 327 data points (Murphy and Romanuk 2014). Climate change has also been documented as a threat to diversity in some groups (Newbold et al. 2020a). However, the colonization of new climatically-suitable areas either by invasive species (Bellard et al. 2018) or species shifting their ranges in response to climate change (Parmesan 2006) can also result in net increases in species richness (Gallardo et al. 2017; Steinbauer et al. 2018). Additionally, the risk for both factors is not distributed evenly across functional groups, taxa or regions: large endotherms, small ectotherms and carnivores are at increased risk (Newbold et al. 2020b), as are taxa found in tropical biomes (Newbold et al. 2020a), perhaps due to higher amounts of intensive disturbance in the tropics (Phillips et al. 2017). Any combination of these contrasting drivers could potentially account for some of the spatial and taxonomic variation in richness change measurements.

Insects, in particular, have declined across a variety of land use gradients (Chisté et al. 2016, Seibold et al. 2019), but particularly due to the conversion of natural land to urban area (Forister et al. 2010; Knop 2016) and the spread and intensification of agricultural landscapes (Raven and Warren 2021). The mechanisms are varied and complex: the application of pesticides (Ewald et al. 2015), the increased input of nitrogen fertilizers (Fox et al. 2014), the loss of obligatory host plants (Koh et al. 2004), the loss of habitat for habitat specialists (Ekroos et al. 2010; Molina-Martinez et al. 2016) and fire suppression (Hahn and Orrock 2015) have all been linked with decreased abundance, species richness and/or beta diversity across insect groups.

Meanwhile, biodiversity has shown mixed responses to forest loss across the globe (Daskalova et al. 2020), with such loss seeming to amplify both increases and decreases in local populations and assemblage species richness through time. For example, forest specialists have been shown to drop out when forest cover declines, but generalists respond positively when the loss of forest cover results in landscape heterogeneity (Estavillo et al. 2013); this seems particularly true in butterflies (Riva et al. 2018).

Importantly, the mass migration of humans to urban areas and widespread agropastoral land abandonment (Levers et al. 2018; Xu et al. 2019) has resulted in the replacement of much traditional cropland and pastures with woodland (Poyatos et al. 2003). This has led to a corresponding diversity increases in some taxonomic groups (Plieninger et al. 2014). In insects, some species such as oak-affiliated butterflies have rebounded in South Korea (Kwon et al. 2021), while grassland-associated Lepidopteran species in Europe have declined (Warren et al. 2021), presumably due to succession (Wölfling et al. 2019).

The effect of these community changes on evolutionary history are less studied, and taxonomic metrics of richness change (or, indeed, lack of change) may not capture the full picture of anthropogenic impacts. Risk is not distributed evenly among clades (Frishkoff et al. 2016). For instance, if closely-related species share similar sensitivities to land use change and climate, the impacts of such changes will be clustered on the phylogeny. Thus, local extinctions (and the corresponding loss of local evolutionary history) may not be random or uniform, but rather clustered on phylogenies. This could potentially result in large swaths of unique evolutionary history being lost, at least at the local level. One metric that can shed light on whether this is happening is Mean Nearest Taxon Distance (MNTD), which calculates the average distance separating every species from its closest relative in an assemblage (Swenson 2014). Because it only looks at nearest neighbor distances, it is sensitive to changes near the tips of the tree (Tucker et al. 2017).

In order to examine the impacts of climate and land-use change on taxonomic and phylogenetic diversity, we focus on Canadian butterflies as a model system. They are a well-suited taxonomic group to examine community responses to anthropogenic change as they have rapid generation times, are sensitive to environmental change, and have been relatively well-documented spatially and temporally. Additionally, Canada represents the northern range limit for all of these species, and provides an ideal opportunity for examining community change in ectotherms as cold-induced climate limits are lifted.

The adverse impacts of intense agriculture on insect diversity have been widely documented and linked to the mechanisms discussed above. As such, we predict the loss of primary or secondary habitat to cropland will result in a loss of species richness, whereas the conversion of cropland to secondary habitat will result in an overall increase in species richness. Insect responses to climate change have been mixed, however, as some species have expanded northwards (Buckley et al. 2012) while others are losing habitat at southern range margins (Settele et al. 2016). As such, we predict no net change in average plot-level species richness in response to climate change.

Meanwhile, the effect of climate change and land-use change on phylogenetic community structure depends on how the relevant traits (climatic and habitat sensitivity) are distributed on the phylogeny. If trait similarity is generally correlated with species relatedness (i.e. there is phylogenetic signal, or niche conservatism), then closely related species should exhibit similar sensitivities to environmental change. Thus, communities that have shifted due to anthropogenic disturbance will comprise more closely related species than communities not experiencing such disturbance (i.e. MNTD should decrease). Conversely, if there is no phylogenetic signal in the relevant traits, and in scenarios where species richness is decreasing (increase in cropland, etc.), species that remain will represent clades scattered throughout the phylogeny (Nowakowski et al. 2018) and communities will increasingly be comprised of more distantly related species. A full schematic of our predictions under each scenario is depicted in Figure 1. Importantly, these predictions are built on the assumption that there is no interspecies competition in these communities: competitive exclusion has the opposite effect on phylogenetic structure than does habitat filtering (Webb et al., 2002). This is likely a reasonable assumption in this group, where drivers such as host plant, habitat and climatic variables have been found to be much more important than interspecific competition in dictating species’ occurrences (e.g. Nakadai et al. 2018).

**Figure 1.**
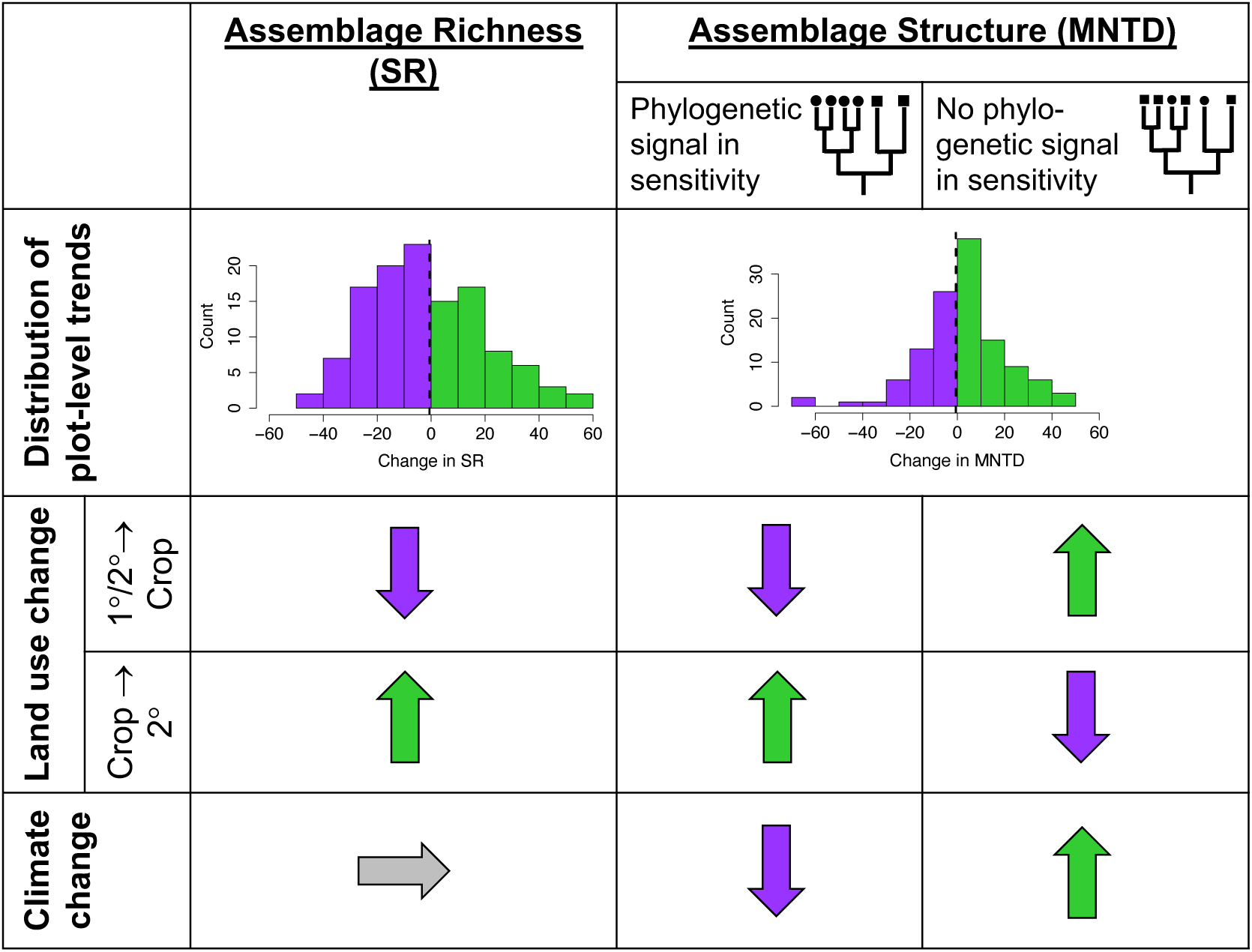
Schematic representation of the environmental metrics used in this study and their hypothesized effect on assemblage richness and assemblage structure. The histograms represent hypothesized plot-level trends through time, where we predict much variation but very little to no overall trend in the average plot-level species richness (SR) or Mean Nearest Taxon Distance (MNTD). Plots that experience increases in SR are illustrated in green on the histogram, whereas decreases are shown in purple. We predict that plot-level changes in SR/MNTD will be influenced by the direction and type of anthropogenic change (land use change or climate change) and the response variable (either SR or MNTD). Predicted directions are illustrated by arrows indicating increases (green), decreases (purple) or no net change (grey) in the response variable. Under MNTD, predictions are further subdivided into whether or not the relevant habitat or climatic niche traits show significant phylogenetic signal.

We test our predictions across 265 species of butterflies and across a wide variety of sites distributed throughout Canada, looking at decadal change from 1940 to 2015. In order to tailor our predictions for our data, we measure phylogenetic signal in climatic niche and habitat-related traits. We then calculate i) species richness change, and ii) change in MNTD, to determine if communities have become significantly more clustered or overdispersed through time. Finally, we consider how climate change and land use change (specifically, agricultural abandonment as well as the loss of natural habitat to agriculture) impact both taxonomic as well as phylogenetic diversity, as measured by species richness and MNTD respectively.

## Methods

### Butterfly Distribution Data

We analyzed approximately 510000 georeferenced and temporally dated butterfly distribution records, spanning 265 species of butterfly that have been observed in Canada. Records were compiled from a variety of sources (see Lewthwaite et al. 2018) and updated with newly submitted eButterfly (see Prudic et al. 2017) records.

Distribution records were amended to follow the latest taxonomic treatment of North American butterflies (Pelham 2014). The average number of records per species across all 96 plots is 322.65 (ranging between 1 and 3873; a full table of record numbers per species can be found in Table S1) The geographical coordinates or identifications for a small proportion of observations (1366) were incorrect, placing species either in regions far beyond their plausible distributions within Canada (Layberry et al. 1998, Brock and Kaufman 2006) or beyond terrestrial limits. As with Lewthwaite et al. 2018, these records were discarded after careful examination.

### Spatial and Community Data

We divided Canada into a system of 10 km x 10 km grid cells, to mimic local assemblages. Similar to Lewthwaite and Mooers (2021), grid cells with fewer than 50 sampled records in both time periods were considered “poorly sampled” and were excluded from maps as well as from the remainder of our analyses. This resulted in 96 cells across Canada that represent the best-sampled quadrats through time (see Figure S4 and S5 for a map of these locations).

Distribution records were divided into decadal bins: 1950-1959, 1960-1969, 1970-1979, 1980-1989, 1990-1999, 2000-2009 and 2010-2015. Opportunistic data (including citizen science), though valuable, has a number of potential biases, including uneven sampling through time (Isaac and Pocock, 2015) and is often not conducive for measuring year-to-year variation. To account for this, we chose to use decadal intervals to represent our species pools as a compromise between uneven sampling effort and a more coarse-scale “before-and-after” comparison (van Swaay 1990). For each grid cell, we summarized the list of species present in each decade, as well as species richness. A total of 265 species were present in these 96 grid cells and within the time period analyzed.

Uneven and incomplete sampling between grid cells and time periods can greatly bias estimates of species richness (Colwell et al. 2012; Chao et al. 2014). As such, we adjusted our species richness estimates with the commonly-used nonparametric asymptotic Chao1 estimator using the ChaoSpecies function in the *SpadeR* package (Chao, Ma, Hsieh & Chiu, 2016). This metric uses the proportion of singletons (species observed only once) to inform the estimate of unseen species (Chao and Chiu, 2016).

With species richness corrections, species identity does not matter. However, species identity is crucial to measuring MNTD. Although methods to adjust PD estimates have recently emerged (Chao et al. 2015; Li 2018), they have not yet been extended to MNTD. Hoever, there is no *a priori* reason why one would expect undersampling within and between quadrats to be phylogenetically biased, and so undersampling may not have a noticeable effect on our MNTD estimate. MNTD values should only be biased by uneven and incomplete sampling if rare species (those that are missed in undersampling) are more likely to be either very evolutionarily distinct (on long branches) or on very short branches. This can be tested.

To test for a relationship between rarity and evolutionary isolation, we used two alternative metrics to quantify a species’ rarity. The first was the number of observations for each species in Canada between 1900-2020s. The second was the number of 10 km x 10 km grid cells occupied by a species in North America across all years. We took two measures of each species’ evolutionary isolation, both being weighted sums of the edge lengths along the path from the root to a tip. The weights of the edge lengths can be distributed evenly among its descendent leaves (i.e. 1/number of species that share that edge; “Fair proportions,” also known as ED), or each edge length can be distributed evenly at each branching point (“Equal splits”) (see, e.g., Wicke and Steel 2020). We calculated isolation via both equal splits and fair proportions using the Canadian butterfly phylogeny (see below) and the *picante* package (Kembel et al. 2010) in R.

We calculated species richness as well as MNTD (see below) for each decadal interval. To measure the change in species richness and MNTD, we looked at the difference between each decade.

### Habitat Affinity and Climatic Niche

In order to test whether sensitivity to land-use change exhibits phylogenetic signal, we examined species’ habitat requirements. Habitat affinities were measured using 4 traits: disturbance affinity, 2) moisture affinity, 3) habitat edge association and 4) canopy preference, all of which are dictated by axes of the environment that would be potentially affected by anthropogenic disturbance. All trait values were obtained via a global database of butterfly traits still under development, which mined field guides and other relevant publications for life history data (Shirey et al. 2022). Trait values were used as ordered categorical predictors, going from one end of the trait spectrum to the other. A list of all categories for each trait and their relative ordering is available in Table S3. Traits were not available for all species used in our analysis. The number of data points for each trait, as well as each trait value are also given in Table S3.

In order to measure the extent to which sensitivity to climate change exhibits phylogenetic signal, we measured each species’ climatic niche width (or the range of climates they have been observed in). We used the same high-resolution (10 km) climate data used in previous chapters, available through the Canadian Forestry Service (CFS; McKenney et al. 2011). This data spans 1901-2010. We chose 5 physiologically relevant climate variables: temperature seasonality (t_s_), maximum temperature of the warmest month (t_w_), minimum temperature of the coldest month (t_c_), precipitation of the wettest quarter (p_w_) and precipitation of the driest quarter (p_d_). These variables were selected because each has empirically supportable, mechanistic links to butterfly physiology and could plausibly limit species’ distributions (Kukal et al. 1991, Crozier 2004, Deutsch et al. 2008, Lewthwaite et al. 2018). For each observation for each species, we extracted the values for each temperature value for the year of the observation. For each species, we then summarized the minimum, maximum and average values across all variables. To calculate the temperature range (t_r_) over which a species has been observed, we took the difference between t_c_ and t_w_.

### Land Use Data

Land use data was obtained from the Land Use Harmonization (LUH2) project (Hurtt et al. 2020), which has estimated the land-use patterns, underlying land-use transitions, and key agricultural management information annually for the time period 850-2100 at 0.25° x 0.25° resolution. Although this is relatively coarse, it is the most comprehensive dataset currently available, both spatially (it has global coverage, including all of Canada) and temporally (it spans the past 10,000 years). This land-use database breaks land use into 12 possible land use ‘states’. For the purposes of this analysis, we grouped them into 5 main land-use types with the following scheme:

1. **Urban**
2. **Cropland**

- C3 annual crops
- C4 annual crops
- C3 perennial crops
- C4 perennial crops
- Nitrogen-fixing crops
3. **Managed Land**

- Pasture
- Rangeland
4. **Primary Land**

- Forested
- Non-forested
5. **Secondary Land**

- Forested
- Non-forested

Units for all states were the fraction of each grid-cell occupied by each land-use state in a given year. In the LUH2 database, primary land is classified as natural vegetation (either forested or non-forested) that has never been impacted by human activities over the past 10,000 years. Meanwhile, secondary land is also classified as natural vegetation (either forested or non-forested), but this vegetation has been previously impacted by human disturbance (again, over the past 10,000 years). This state can range from very young vegetation recovering from a recent human disturbance, to vegetation that is very mature, recovering from a much older human disturbance.

The LUH2 database also quantifies rates of transitions between land-use states, either at a decadal scale (prior to 2000) or a yearly scale (post-2000). The land-use transitions give the fraction of each grid-cell that transitions one from land-use state to another in a given decade/year, and is therefore a rate of change. There are no transitions from any land use state to primary (e.g. secondary land can never return to a “primary” state, even if it is very mature).

Looking at the absolute amount of each land-use state through time per grid cell can only yield so much information. For example, it would indicate if there was an increase in secondary habitat within a grid cell through time but it would not specify what that habitat was being converted from. If it were a conversion from primary land to secondary land, this may yield very different outcomes and predictions than if it were a conversion from cropland to secondary land. Therefore, we decided to focus on specific transitions between habitat types through time, rather than relative amounts of each land use state.

We were specifically interested in 3 types of land use transitions that we had strong predictions for: i. conversion of primary habitat to cropland; ii. conversion of secondary habitat to cropland (both of which represent the loss of natural habitat to agriculture); and iii. conversion of cropland to secondary habitat (as a metric of agricultural abandonment).

To convert the land use transition files from the NetCDF file format to rasters, we used the *ncdf4* and *raster* packages in R (Pierce, 2021; Hijmans, 2020). For each year, we then stacked the rasters for each transition type we were interested in (for example, for conversion of cropland to secondary, we stacked the transitions from all the cropland subcategories (C3 annual, C4 annual, etc.) to secondary subcategories (forested, non-forested)). This resulted in a sum of yearly/decadal transitions for each of the 3 transition types in each cell. We then calculated the mean transition rate for each of the transition types within each of the 96 quadrats for each decade.

There is evidence for a lag in response times in invertebrates to land use change of around 7 years (Daskalova et al. 2020). Previous work on pollinator responses to habitat change has incorporated a metric of historic land use change in order to account for this lag (see, e.g. Aguirre-Gutiérrez et al. 2015). Many have found a significant effect of historical land use on butterfly species or communities (Casner et al. 2014; Cusser et al. 2015; Cusser et al. 2018). Therefore, we decided to incorporate a metric of historical land use change in our models by including a term for the land use conversion rate for the previous decade, as well as the contemporary period. We did this for all 3 conversion types (primary to cropland, secondary to cropland, and cropland to secondary).

### Quantifying Climate Change

To quantify the rate of climate change experienced within each grid cell in each decade (e.g. the 1950s), we first extracted the mean temperature for each year within that decade (e.g. 1950, 1951, 1952, etc.) from the annual CFS climate data discussed above. We then calculated the difference between each year to get the year-to-year variation (e.g. the difference between 1951 and 1950, the difference between 1952 and 1951, etc.). Finally, we took the average of all the year-to-year differences within a cell. This gave us the mean yearly temperature change within a cell across the decade, or the decadal rate of change. For example, for cells that gradually increased each year in temperature, the average year-to-year difference would be a small increase in temperature. However, if the temperature within a cell fluctuated year-to-year but with no directional change, the average difference in mean temperature would be close to zero.

As the climate data from CFS only extends until 2010, we repeated the same workflow as above for 2010-2015 using data from the Climate Research Unit (Harris et al. 2020), which is a global dataset, though it is not as high resolution (0.5° x 0.5°, or roughly 55 kilometers square) as the CFS data (10km square).

### Phylogenetic Analyses

We constructed a phylogeny of all Canadian butterfly species using eight mitochondrial and protein-coding nuclear genes, totalling 7178 nucleotides with gaps.

The phylogeny was generated using BEAST ver 1.10. (Suchard et al. 2018), and was run for 125 million generations with a 25% burn-in fraction. Species that were missing genetic data were constrained in their placement in the tree using taxonomy. The phylogeny was dated using four fossil calibrations and the maximum height of the root of the tree could not predate the origin of the angiosperms at 183 MYA (Bell et al. 2010), based on the assumption that all but the earliest Lepidoptera were constrained by the diversification of angiosperms (Condamine et al. 2012). A full description of methods used to generate the phylogeny can be found in the Supporting Information.

We ran all analyses across a distribution of 1000 possible tree topologies that were randomly sampled from the posterior distribution.

We calculated the phylogenetic signal for each habitat trait and climatic variable using the *phytools* package in R (Revell, 2012). For each trait, we calculated Pagel’s lambda, where a value close to zero indicates phylogenetic independence, whereas a value of one indicates that species’ traits are distributed as expected under Brownian Motion (Münkemüller et al. 2012).

We used the *picante* package in R (Kembel et al. 2010) to measure MNTD for Time 1 and Time 2, as well as for each decade. For each, we took the average value across all 1000 candidate topologies for later analyses.

MNTD is not completely independent from species richness, and the default expectation is that it decreases with increased species richness (Cadotte et al. 2010). We verified that this was the case with our data. Adjusted (Chao1) species richness estimates within the 96 quadrats used in this study ranges from 8 species to approximately 103 species. Thus, we randomly pruned our phylogeny of all Canadian butterflies in order to obtain a distribution of trees that ranged from 8 to 100 species. We then extracted MNTD from each tree and compared that to the number of tips for that tree (species richness). Indeed, we do find a negative relationship between MNTD and species richness, particularly in communities with low species richness (Figure S2). However, when species are added or removed from communities, this relationship is not always straightforward: when close relatives are added to a community, MNTD may decrease (Baiser et al. 2018), whereas an introduction of an exotic and distantly-related species could result in an increase in MNTD (McGrannachan et al 2020). Therefore, any change in MNTD must be interpreted alongside change in species richness. As such, we controlled for species richness change when examining change in MNTD (see below).

### Statistical Analyses

All final statistical analyses were conducted in a Bayesian framework, using the R package *brms* (Bürkner 2017) in R v3.6.0. The *brms* package allows the user to fit generalized multivariate and multilevel models in a Bayesian framework using Stan (a C++ program) (Carpenter et al. 2017). We chose to use a Bayesian approach in order to assess the level of uncertainty in our model.

We scaled and centered all predictors so that effects are comparable between predictors and the units of the regression coefficients are the same. Each was included in the model as a fixed effect with non-informative priors.

We constructed two separate models: one to model change in species richness, and one for change in MNTD. For each model, we used decades as the temporal interval, and measured the change in either the predictor or response variable in each of the 96 quadrats, so that each data point is a grid cell in each decade. Each model included 6 metrics of land-use change as fixed effects: the rate of contemporary and historic conversion for each of the 3 types of transition (primary to cropland, secondary to cropland, and cropland to secondary) within each decade. We also included a fixed effect for the mean temperature change a grid cell experienced to account for climate change (see above). Finally, grid cell identity was added as a random effect to account for any variability between grid cells that were unaccounted for by the fixed effects. We included change in species richness as a fixed effect in the MNTD model. We constructed 4 Markov chains of 4000 iterations each, including a warm-up phase of 1000 iterations, and used an adapt_delta value of 0.99 to reduce the number of divergent transitions. Specific model structures and priors can be found in the Supporting Information.

## Results

### Phylogenetic tree and signal

All of the major and early splits in our tree were strongly supported, as evidenced by the high bootstrap values (Figure S1). However, there were some low-support nodes near the tips, particularly in groups that were data-deficient. Our estimates of crown ages for the 5 major families and the crown of butterflies were largely in line with what has been found in other studies. We found little evidence of phylogenetic signal in both of our habitat and climate-related traits (Table 1).

**Table 1.** Phylogenetic signal of habitat affinities and climatic niche across butterfly species.

| Trait | # of species | Lambda | logL(lambda) |
| --- | --- | --- | --- |
| Disturbance Affinity | 205 | 0.44 | -384.61 |
| Canopy Cover | 261 | 0.18 | -318.00 |
| Edge Association | 249 | 0.05 | -427.73 |
| Moisture Affinity | 256 | 0.30 | -470.30 |
| Temperature Range | 298 | 0.08 | -1654 |
| Min. Temp (coldest month) | 298 | 0.4 | -1748 |
| Max. Temp (warmest month) | 298 | 0.5 | -1563 |

### Evolutionary distinctiveness vs rarity

We found no evidence of a relationship between a species’ Canadian isolation score (measured as either equal splits or fair proportions) and their rarity (measure as either the # of observations, or their range size; Figure S3). As such, there is no evidence that undersampling in our study would significantly bias our MNTD estimates by missing species on particularly long or short branches on the phylogeny.

### Land use change

Since 1945, the most significant land use changes across Canada have been a loss of primary habitat (forest and non-forest) and an increase in secondary habitat (forest and non-forest) (see Figure 2), the former likely driven by a combination of logging, fire, forest disease and resource extraction (Hansen et al. 2013; Potapov et al. 2017) and the latter likely due to recent global trends of agropastoral land abandonment (Levers et al. 2018; Xu, Deng, Guo & Liu, 2019) including within Canada (Potapov et al. 2022).

**Figure 2.**
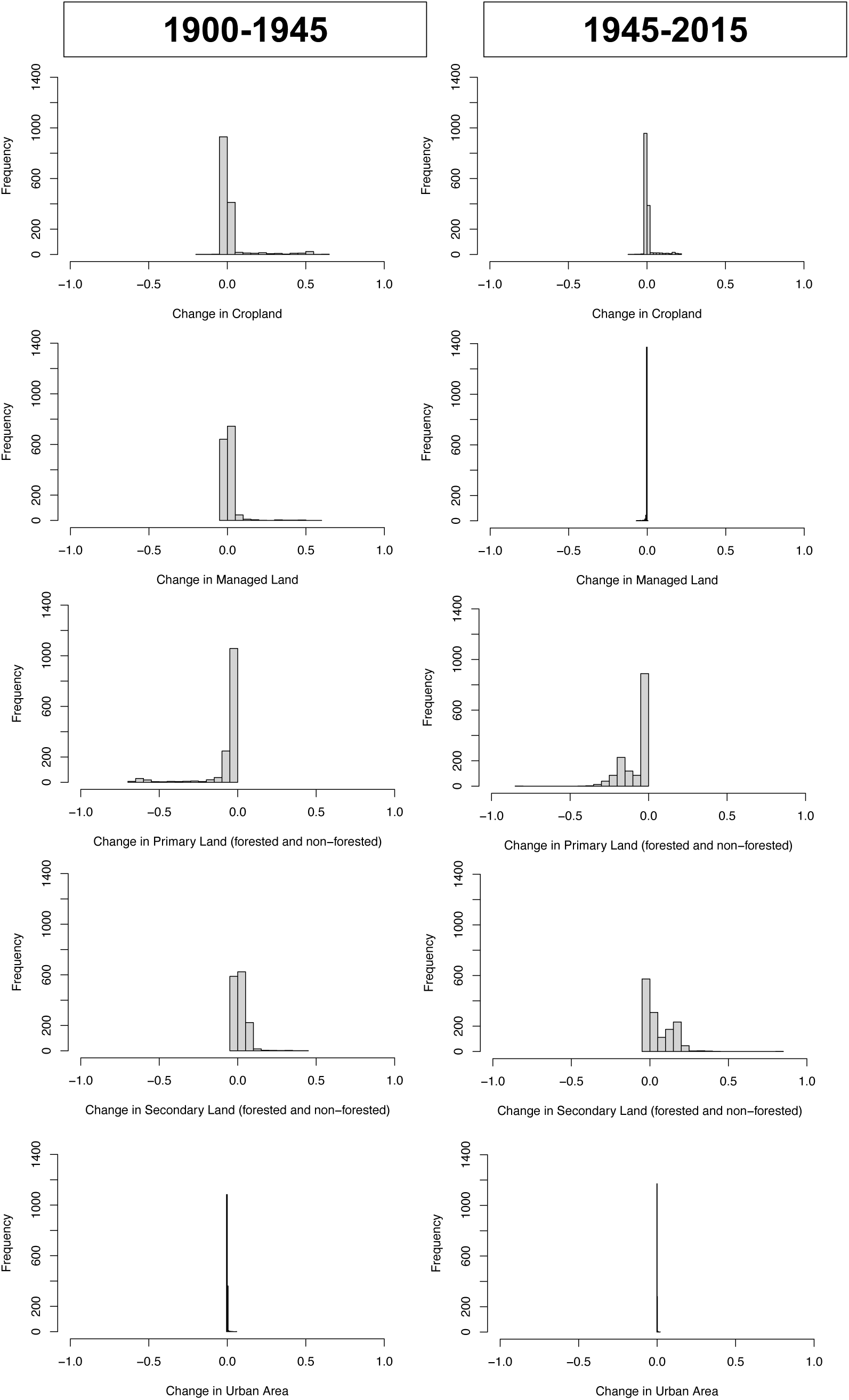
Histograms of land use change across Canada within two time periods: 1900-1945 and 1945-2015. Land use data was divided into 5 main categories: Cropland (including c3ann, c4ann, c3per, c4per and c3nfx), Managed (including rangeland and pasture), Primary (including forest and non-forest), Secondary (including forest and non-forest), and Urban. Summary statistics were obtained by dividing Canada into a series of 100 km x 100 km grid cells and calculating the proportion of each land use category within it. Change in land use measured as the differences in proportions of each land use category within a grid cell between the two time periods (1900-1945 vs. 1945-2015). n = 1518 quadrats.

Comparatively little area has been converted to urban landscapes across both time periods. This is consistent with urban areas in North America (and particularly Canada) being relatively localized and occupying a relatively small proportion of total land area (Seto et al. 2012). Additionally, the conversion of land to or from managed land as well as cropland has slowed substantially since 1945.

### Species richness and MNTD patterns in Canadian butterflies

MNTD values across Canada are lowest in the southern portions of the country, and highest in the North (Figure S5A). Species richness also tends to be highest in southern Canada (Figure S4A). However, there is little spatial variation in MNTD in the southern half of the country, whereas there is much more variation in species richness. This is likely because above a certain richness threshold, any addition or loss of a species has a minor impact on MNTD. This is driven by two factors. The first is that MNTD is calculated as an average and is less likely to vary by large amounts when species richness is high. The second is that when species richness is high, the chances are much higher that an added or lost species already has a close relative in the phylogeny. However, when species richness drops below a certain point (a point passed in the North), it is much less likely that any species added/lost has a close relative already present. The species that are present are more dispersed on the phylogeny, and so any variation becomes more significant.

Overall, there were 419 measurements of species richness change between all decade and quadrat combinations (out of a potential 672). The Chao1 species richness adjustor cannot adjust estimates when there are only singletons (species only observed once within a plot) across all quadrats in a decade. Therefore, we were unable to calculate change between decades where one decade had no available Chao1 estimate. There were 464 measurements of MNTD change. As MNTD was based on raw species richness data (not the Chao1 estimates) and it is a pairwise metric, we could not calculate MNTD in situations where there was only 1 species observed in a quadrat. Overall, there were 408 quadrat and decade combinations that had estimates for both change in species richness and change in MNTD.

Average species richness within these 96 quadrats is 39.47 species (SD=31.10). Across quadrats, the average change in species richness between decades is an increase of 3.95 species per decade (SD=29.84). The corresponding change in average MNTD was a decrease of 3.85 million years (SD=47.57; Figure 3A). This is a statistically insignificant change in MNTD (95% CI -8.19 – 0.48, unpaired two-tailed t-test, t=-1.74, df = 463, p=0.08).

**Figure 3.**
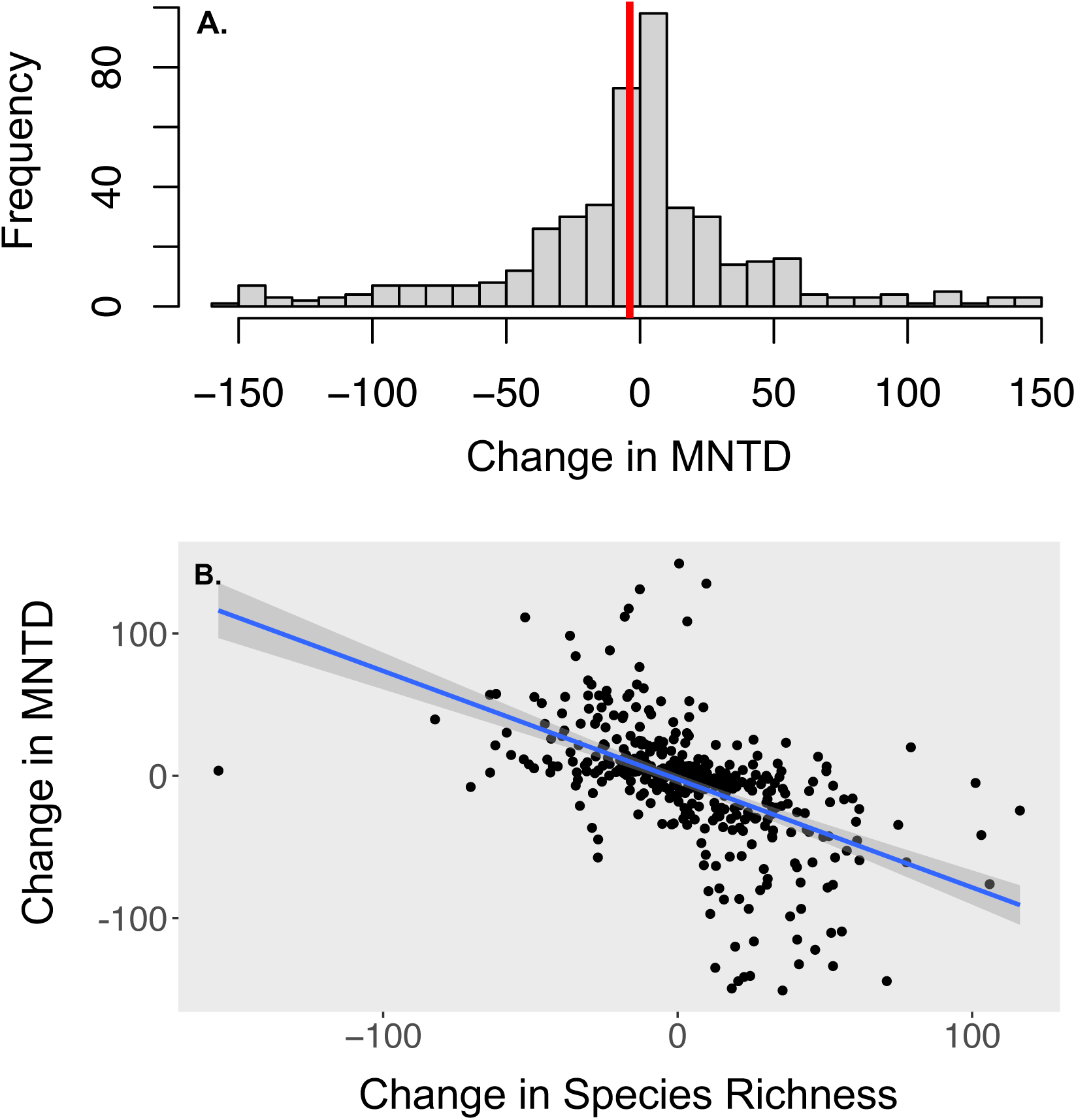
(A) Histogram of the observed change in Mean Nearest Taxon Distance (MNTD) between decades; and (B) the relationship between the change in MNTD (averaged across 1000 candidate topologies) and the change in species richness within a quadrat between decades. Red line in (A) represents the observed average. A linear regression and the 95% confidence interval of the predicted values are overlaid in (B). n = 408 decade + quadrat combinations.

There is a significant negative relationship between change in species richness and change in MNTD (linear regression b=-0.76, SE=0.06, t= -12.54, p<0.001, R^2^ = 0.28, F_1,407_=157.3; Figure 3B). Thus, as an assemblage gains species through time, the distance between co-occurring species decreases, meaning that species on average are now more closely-related to their closest relative in the assemblage. And, if an assemblage loses species, the distance increases i.e. species are less-closely-related to their closest relative in the assemblage).

There was no obvious pattern in where the largest increases and decreases in MNTD between decades occurred in Canada (Figure S5B), though many of the largest decreases occurred within the same quadrat but in separate decades. The magnitude and direction of species richness and MNTD change varies between decades (Table S2, Figure S6). The 1970s experienced the largest increase in species richness, whereas the 1990s saw the largest decrease in species richness. MNTD generally declined between decades, except for an increase in the 1980s and 1990s.

None of the climatic or land-use predictors were significantly associated with change in species richness or MNTD between decades (Figure 4). However, mean temperature change and the historical conversion of secondary habitat to cropland had relatively large and negative impacts on species richness within the relevant decade (Figure 4A), whereas recent agricultural abandonment lead to increases in species richness. Meanwhile the contemporary loss of primary habitat to cropland saw an increase in MNTD (though not significantly) within the decade, where species within an assemblage were more distantly related to each other (Figure 4B), while contemporary agricultural abandonment lead to a decrease in MNTD.

**Figure 4.**
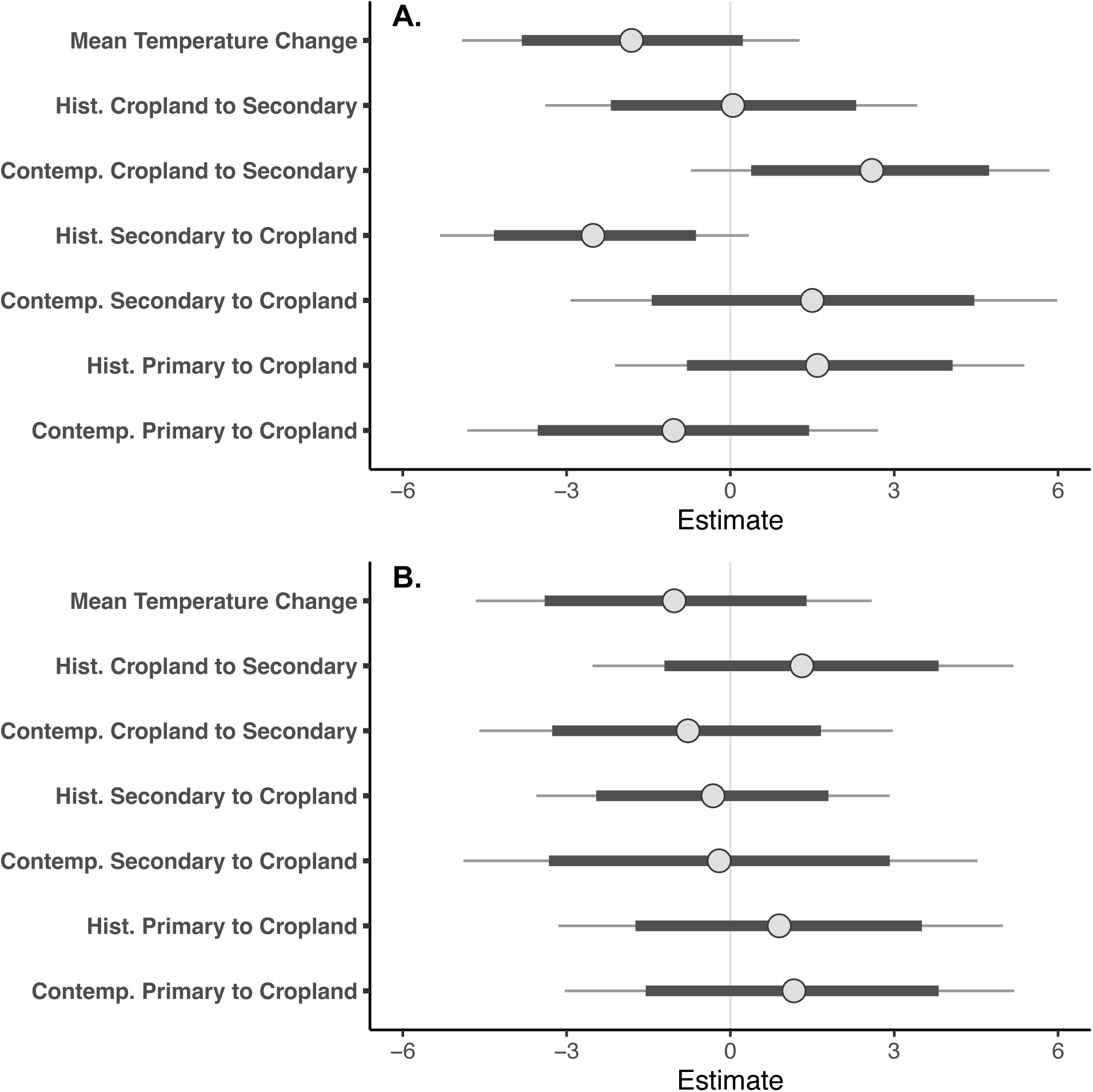
Posterior distributions of estimates for each of the land-use and climate transition rates examined and their relationship to (A) the change in species richness; and (B) the change in Mean Nearest Taxon Distance (MNTD; averaged over 1000 candidate topologies, and controlling for species richness change). The thin lines for each distribution represents the 95% confidence intervals around the estimate; the thicker lines represents the 80% confidence interval. n = 408 decade and quadrat combinations.

## Discussion

There is evidence that contemporary climate change has both positively and negatively affected different insect groups (Halsch et al. 2021), potentially leading to the “no net change” pattern, as documented in Lewthwaite and Mooers (2021). Similarly, the effects of land use change have been mixed. The most geographically and taxonomically representative models to date show precipitous declines in both species richness and abundance in intensively-used landscapes (Newbold et al. 2015). Likewise, when areas of traditional low-intensity land-use are abandoned for agricultural intensification, it is widely accepted that this results in a decrease in insect diversity (Uchida and Ushimaru, 2014; Uchida et al. 2016), likely as a result of pesticide use, homogenization of landscapes and resources and habitat fragmentation (Habel et al. 2019). However, when these are abandoned in turn as a result of human migration to cities, this can cause declines in open-habitat butterfly species (Scherer et al. 2021), and simultaneously, increases in forest butterflies (Herrando et al. 2016).

Compared to pre-1945 levels, there has been relatively little conversion of land to or from cropland in Canada (Figure 2). As such, although secondary habitat is increasing in Canada since 1945, this is largely not as a result of a loss in cropland. Similarly, Canada is no longer losing much secondary habitat to cropland. However, the small amounts of both types of conversion that do occur seem to have large effects on a quadrat’s species richness, both negative and positive (Figure 4). While the conversion of primary and secondary habitat to cropland has resulted in mixed trends in richness depending on how much time has passed, the abandonment of cropland and subsequent succession to secondary habitat has resulted in immediate increases in species richness that eventually taper off (Figure 4A). One of the most striking trends is a decrease in species richness in quadrats that have experienced the largest temperature changes through time. Although a warming climate might initially seem beneficial for this group as thermal limits are lifted at northern range boundaries, Lewthwaite et al. (2018) found that many of these species are not colonizing newly climatically-suitable area. Similar lags have been observed in British butterflies (Menéndez et al. 2006). Extreme events linked to climate change have also been linked to population declines and community structure changes (Halsch et al. 2021), either directly (ex. via daytime heat stress, or through increased precipitation during critical overwintering stages) or indirectly (ex. through their host plant). Furthermore, a change in site-level maximum temperature has been shown to be the climate variable most linked with local extinctions in insects (Román-Palacios and Wiens, 2020). A potential depletion of climate specialists combined with the lack of replacement by other species (who are lagging behind the pace of climate change) may be responsible for the large species richness declines that we observe in cells that are warming the fastest.

Meanwhile, the effect of agricultural intensification depended on the starting habitat and changed with the amount of time since conversion, which could potentially indicate a lag effect. When it was secondary habitat, the initial conversion to cropland resulted in an increase in species richness. However, this turned into a negative effect with time. Secondary habitat is previously disturbed area, which are likely composed of species that prefer open or mixed canopies (which are the majority of species; see Table S3) and are disturbance-affiliated. As such, the further clearing of canopy perhaps initially resulted in local colonizations from nearby areas, but richness declined in later years as agriculture intensified and less-resilient species dropped out. However, we see the opposite trend when the starting habitat is primary; we see an initial decrease in species richness as it is converted to agriculture. Untouched primary habitat may be more likely to be composed of closed canopy and disturbance-avoidant species, which are initially lost in the conversion to agriculture. However, because there are many more open/mixed canopy and disturbance-affiliated species than the converse, potential later colonization by the most agriculturally-resilient species could result in the eventually rebound in richness levels that we see.

Indeed, if traits mediate much of the varied responses we see in our own study as well as the literature, and if species that share more evolutionary history are more likely to share similar trait spaces (Tucker et al. 2018), then one might expect species to be non-randomly filtered out of communities in the face of both land use change (Egorov et al. 2014) and climate change (Davis et al. 2010). A clustered loss of species on the tree may reduce functional diversity and potentially reduce ecosystem services (Grab et al. 2019). Our empirical results showed that although there is much variation in the change in MNTD within a community, mean quadrat-level change is indistinguishable from zero (Figure 3A).

We also found a significantly negative relationship between change in species richness and change in MNTD within an assemblage, though the modest R^2^ (0.28) illustrates lots of variation in this relationship, particularly around the mean (3.85 MY; Figure 3B). An increase in species richness within an assemblage rarely resulted in an increase in MNTD, which likely indicates that colonizing species (such as introduced species) were usually close relatives of members already within that quadrat and were not adding much novel phylogenetic information to an assemblage. Similarly, a decrease in species richness rarely resulted in a decrease in MNTD, indicating that as species drop out of assemblages, the remaining species are less closely related (as expected). Species that were locally extirpated were likely to be distributed across the tree and not highly evolutionary distinct, as the loss of highly evolutionarily isolated species would actually decrease MNTD in a plot, as remaining species would now be more closely-related than they previously were. Additionally, if losses were clustered on the phylogeny, a decrease in species richness would result in a decrease in MNTD, which was rarely the case in our assemblages.

Though there was no net change in MNTD (and MNTD exhibited far less variability in response than species richness to land use or climate change; Figure 4B), there was some variation in this response between assemblages. The most notable (though still statistically insignificant) changes were an increase in MNTD in response to the contemporary conversion of primary habitat to cropland, and a decrease in MNTD in response to a contemporary conversion of cropland to secondary habitat. Most of the traits have very low phylogenetic signal (Table 1), which would in theory indicate that closely-related species are not more likely to share sensitivities to environmental change. As such, many of the observed changes in MNTD in response to land-use change are consistent with our predictions (see Figure 1), though we did not see all of the expected responses to land use change that we predicted.

Contrary to our predictions, mean temperature change within a quadrat resulted in a steep decline in species richness, and a slight decrease in MNTD. (Figure 4). Therefore, the species that remain tend to be more closely-related than expected, indicating that the impacts of climate change may be clustered on the phylogeny.

We see positive effects of agricultural abandonment, with a relatively large increase in species richness in the years immediately following this. Even after controlling for this increase in species richness, MNTD decreased initially. However, MNTD later increased with no change in species richness, indicating a re-structuring of these communities.

Our results are also consistent with trends in the literature with regard to phylogenetic restructuring in response to agricultural change (Kusuma et al. 2018). Although species richness declined in quadrats where there was a recent loss of primary habitat to cropland, the mean distance between species increased. This indicates that while there are typically fewer species in converted habitats, these species represent clades scattered throughout the phylogeny rather than clustered in groups (Nowakowski et al. 2018). Conversely, a gain in species richness in cropland that was recently abandoned corresponded to a decrease in mean distance between community members. Species that were added tended to be close relatives of members that were already present.

Interestingly, this would also suggest that relevant traits are potentially being conserved – but not the ones we measured. The species arriving into assemblages tend to be close relatives to those that persisted following habitat change, suggesting that habitat affinity does show phylogenetic clumping. However, we found no significant phylogenetic signal in the traits that we measured. The tree itself may therefore be a better representation of ecological affinity than our own attempts to derive these metrics – suggesting that the tree is integrating ecology, and that preserving phylogenetic diversity may save much more than we realize.

There are some additional considerations in our analysis. Although the increase in secondary habitat is likely driven by a concurrent loss of primary habitat, we did not assess the biotic impacts of primary to secondary habitat conversion, as any predictions were not straightforward or generalizable across all species. Some woodland butterfly species would have likely declined (van Swaay et al. 2006), whereas grassland species may have benefitted (Viljur and Teder, 2016), and so responses would be species-specific.

Additionally, this study looks at the relative change but not the absolute amount of each category type within a grid cell. For example, cropland may have only slightly increased within a cell through time, but that may not mean much if it was already mostly cropland to begin with.

Finally, much of the loss of primary habitat and increase in cropland in Canada occurred prior to 1945 (Figure 2). It is possible that we have missed much of the resulting species and phylogenetic change that resulted prior to 1945. Much of the loss in diversity and homogenization of communities may have already occurred by the 1950s and so our study may only be examining the tail end of that change. Any increases in species richness or phylogenetic diversity from recent agricultural abandonment may not make up for that initial loss.

Initially, these findings would imply some good news for biodiversity conservation in Canada, as we found that modern agricultural abandonment is associated with net species richness increases in our quadrats. However, the full picture is more nuanced. Firstly, when species are initially lost in the conversion to cropland, they tend to be scattered throughout the phylogeny. However, when areas are recolonized by species following agricultural abandonment, they tend to be close relatives of the species already present. Secondly, because the actual amount of this land use conversion type is so small in these quadrats, it is unlikely to have or to have had a large-scale impact on Canadian biodiversity.

Therefore, as we have noted in previous work (Lewthwaite and Mooers, 2021), by only examining one axis of biodiversity in a community, such as species richness, one may miss other diversity changes that are simultaneously occurring. We also find that much of the variation we see in Canadian butterfly diversity trends through time is likely influenced by past anthropogenic change, indicating that shifts in community structure are likely to continue into the future.

## Acknowledgements

Thanks to members of the Crawford Lab at SFU for ongoing input to this manuscript. J.M.M.L. was supported by the Natural Sciences and Engineering Research Council of Canada (NSERC) through a doctoral scholarship (PGS-D) and A.Ø.M. through a Discovery and an Accelerator Grant. We thank eButterfly and all of their affiliated citizen scientists for collecting, reporting and vetting butterfly observation data used in this publication. We also thank Heather Kharouba for compiling some of the data used in this manuscript, Ross Layberry for help with georeferencing, as well as Guy Baillargeon, Cris Guppy, Norbert Kondla, Ross Layberry, Karen Needham and the Toronto Entomological Association for providing observation data.

