## Supporting Information for "75 years of anthropogenic change and its impact on Canadian butterfly taxonomic and phylogenetic diversity"

### This file includes:

Figs. S1 to S7

Tables S1 to S11

Supplementary Materials and Methods

Part A: Choice of priors for Bayesian analysis of quadrat-level responses to land-use and climate change

Part B: Habitat traits

Part C: Phylogeny construction

References

### Supplementary Results

**Table S1:** Number of records per species across all 96 plots from 1940-2015. n=265 species.

| Species | # of Records | Species | # of Records |
|---|---|---|---|
| Abaeis nicippe | 3 | Chlosyne hoffmanni | 15 |
| Aglais milberti | 676 | Chlosyne nycteis | 246 |
| Agriades glandon | 213 | Chlosyne palla | 209 |
| Agriades optilete | 47 | Coenonympha tullia | 3308 |
| Amblyscirtes hegon | 41 | Colias alexandra | 85 |
| Amblyscirtes vialis | 311 | Colias canadensis | 53 |
| Anatrytone logan | 265 | Colias christina | 152 |
| Ancyloxypha numitor | 515 | Colias eurytheme | 1171 |
| Anthocharis sara | 104 | Colias gigantea | 64 |
| Anthocharis stella | 376 | Colias hecla | 56 |
| Asterocampa celtis | 347 | Colias interior | 238 |
| Asterocampa clyton | 248 | Colias meadii | 124 |
| Atalopedes campestris | 26 | Colias nastes | 120 |
| Atrytonopsis hianna | 79 | Colias occidentalis | 87 |
| Battus philenor | 83 | Colias palaeno | 132 |
| Boloria alaskensis | 47 | Colias pelidne | 39 |
| Boloria alberta | 29 | Colias philodice | 3129 |
| Boloria astarte | 106 | Cupido amyntula | 395 |
| Boloria bellona | 539 | Cupido comyntas | 1084 |
| Boloria chariclea | 318 | Danaus plexippus | 2799 |
| Boloria epithore | 85 | Echinargus isola | 1 |
| Boloria eunomia | 176 | Epargyreus clarus | 627 |
| Boloria freija | 308 | Erebia disa | 6 |
| Boloria frigga | 119 | Erebia discoidalis | 151 |
| Boloria improba | 35 | Erebia epipsodea | 525 |
| Boloria polaris | 58 | Erebia fasciata | 37 |
| Boloria selene | 368 | Erebia lafontainei | 17 |
| Callophrys affinis | 41 | Erebia mackinleyensis | 27 |
| Callophrys augustinus | 589 | Erebia magdalena | 29 |
| Callophrys eryphon | 199 | Erebia mancinus | 108 |
| Callophrys gryneus | 280 | Erebia occulta | 8 |
| Callophrys henrici | 305 | Erebia pawlowskii | 86 |
| Callophrys irus | 34 | Erebia rossii | 74 |
| Callophrys johnsoni | 7 | Erebia vidleri | 51 |
| Callophrys lanoraieensis | 9 | Erebia youngi | 29 |
| Callophrys mossii | 175 | Erora laeta | 5 |
| Callophrys niphon | 397 | Erynnis afranius | 12 |
| Callophrys polios | 300 | Erynnis baptisiae | 314 |
| Callophrys sheridanii | 122 | Erynnis brizo | 134 |
| Callophrys spinetorum | 27 | Erynnis funeralis | 10 |
| Carterocephalus palaemon | 527 | Erynnis horatius | 54 |
| Celastrina echo | 362 | Erynnis icelus | 748 |
| Celastrina ladon | 184 | Erynnis juvenalis | 717 |
| Celastrina lucia | 1488 | Erynnis lucilius | 300 |
| Celastrina neglecta | 937 | Erynnis martialis | 94 |
| Celastrina serotina | 33 | Erynnis pacuvius | 86 |
| Cercyonis oetus | 112 | Erynnis persius | 458 |

| | | | |
|---|---|---|---|
| Cercyonis pegala | 1411 | Erynnis propertius | 180 |
| Cercyonis sthenele | 27 | Euchloe ausonides | 354 |
| Chlosyne damoetas | 13 | Euchloe creusa | 78 |
| Chlosyne gorgone | 41 | Euchloe lotta | 77 |
| Chlosyne harrisii | 179 | Euchloe naina | 15 |
| Euchloe olympia | 216 | Nymphalis antiopa | 2312 |
| Euphilotes battoides | 113 | Nymphalis californica | 259 |
| Euphydryas anicia | 579 | Nymphalis l-album | 556 |
| Euphydryas chalcedona | 28 | Oarisma garita | 88 |
| Euphydryas editha | 221 | Ochlodes sylvanoides | 555 |
| Euphydryas gillettii | 25 | Oeneis alberta | 44 |
| Euphydryas phaeton | 272 | Oeneis alpina | 17 |
| Euphyes bimacula | 62 | Oeneis bore | 151 |
| Euphyes conspicua | 32 | Oeneis chryxus | 275 |
| Euphyes dion | 158 | Oeneis jutta | 182 |
| Euphyes dukesi | 64 | Oeneis macounii | 44 |
| Euphyes vestris | 883 | Oeneis melissa | 86 |
| Euptoieta claudia | 168 | Oeneis nevadensis | 58 |
| Eurytides marcellus | 18 | Oeneis philipi | 19 |
| Feniseca tarquinius | 214 | Oeneis polixenes | 78 |
| Glaucopsyche lygdamus | 1596 | Oeneis uhleri | 64 |
| Glaucopsyche piasus | 106 | Panoquina ocola | 12 |
| Hesperia assiniboia | 46 | Papilio brevicauda | 19 |
| Hesperia colorado | 152 | Papilio canadensis | 840 |
| Hesperia comma | 112 | Papilio cresphontes | 639 |
| Hesperia juba | 101 | Papilio eurymedon | 247 |
| Hesperia leonardus | 168 | Papilio glaucus | 810 |
| Hesperia nevada | 23 | Papilio indra | 5 |
| Hesperia sassacus | 184 | Papilio machaon | 182 |
| Hylephila phyleus | 282 | Papilio multicaudata | 91 |
| Icaricia icarioides | 516 | Papilio polyxenes | 1096 |
| Icaricia lupini | 120 | Papilio rutulus | 380 |
| Icaricia saepiolus | 463 | Papilio troilus | 458 |
| Icaricia shasta | 1 | Papilio zelicaon | 175 |
| Junonia coenia | 390 | Parnassius clodius | 88 |
| Leptotes marina | 11 | Parnassius eversmanni | 48 |
| Lerema accius | 1 | Parnassius phobeus | 12 |
| Lethe anthedon | 562 | Parnassius smintheus | 401 |
| Lethe appalachia | 170 | Parrhasius m album | 9 |
| Lethe eurydice | 464 | Phoebis sennae | 34 |
| Libytheana carinenta | 318 | Pholisora catullus | 246 |
| Limenitis archippus | 1085 | Phyciodes batesii | 210 |
| Limenitis arthemis | 1293 | Phyciodes cocyta | 1943 |
| Limenitis lorquini | 430 | Phyciodes mylitta | 286 |
| Lycaena cupreus | 45 | Phyciodes pallida | 24 |
| Lycaena dione | 48 | Phyciodes pulchella | 276 |
| Lycaena dorcas | 145 | Phyciodes tharos | 516 |
| Lycaena epixanthe | 111 | Pieris angelika | 106 |
| Lycaena helloides | 470 | Pieris marginalis | 172 |
| Lycaena heteronea | 144 | Pieris oleracea | 545 |
| Lycaena hyllus | 462 | Pieris rapae | 3873 |
| Lycaena mariposa | 80 | Pieris virginiensis | 161 |
| Lycaena nivalis | 42 | Plebejus idas | 408 |

| | | | |
|---|---|---|---|
| Lycaena phlaeas | 501 | Plebejus melissa | 396 |
| Megisto cymela | 1083 | Poanes hobomok | 1052 |
| Nathalis iole | 3 | Poanes massasoit | 20 |
| Neominois ridingsii | 3 | Poanes viator | 96 |
| Neophasia menapia | 184 | Polites draco | 33 |
| Polites mystic | 612 | Wallengrenia egeremet | 483 |
| Polites origenes | 220 | Zerene cesonia | 7 |
| Polites peckius | 454 | | |
| Polites sabuleti | 36 | | |
| Polites sonora | 4 | | |
| Polites themistocles | 575 | | |
| Polygonia comma | 843 | | |
| Polygonia faunus | 289 | | |
| Polygonia gracilis | 158 | | |
| Polygonia interrogationis | 937 | | |
| Polygonia oreas | 15 | | |
| Polygonia progne | 365 | | |
| Polygonia satyrus | 218 | | |
| Polyommatus icarus | 5 | | |
| Pompeius verna | 129 | | |
| Pontia beckerii | 59 | | |
| Pontia occidentalis | 382 | | |
| Pontia protodice | 53 | | |
| Pontia sisymbrii | 78 | | |
| Pyrgus centaureae | 125 | | |
| Pyrgus communis | 79 | | |
| Pyrgus ruralis | 167 | | |
| Pyrisitia lisa | 174 | | |
| Satyrium acadica | 415 | | |
| Satyrium behrii | 26 | | |
| Satyrium calanus | 498 | | |
| Satyrium californica | 49 | | |
| Satyrium caryaevorous | 153 | | |
| Satyrium edwardsii | 172 | | |
| Satyrium liparops | 353 | | |
| Satyrium saepium | 59 | | |
| Satyrium semiluna | 14 | | |
| Satyrium sylvinus | 43 | | |
| Satyrium titus | 347 | | |
| Speyeria aphrodite | 301 | | |
| Speyeria atlantis | 240 | | |
| Speyeria callippe | 124 | | |
| Speyeria cybele | 1083 | | |
| Speyeria edwardsii | 14 | | |
| Speyeria hesperis | 137 | | |
| Speyeria hydaspe | 204 | | |
| Speyeria idalia | 5 | | |
| Speyeria mormonia | 330 | | |
| Speyeria zerene | 345 | | |
| Staphylus hayhurstii | 7 | | |
| Strymon melinus | 607 | | |
| Thorybes bathyllus | 19 | | |
| Thorybes pylades | 570 | | |

| | |
|---|---|
| Thymelicus lineola | 1131 |
| Vanessa annabella | 67 |
| Vanessa atalanta | 1979 |
| Vanessa cardui | 832 |
| Vanessa virginiensis | 1007 |

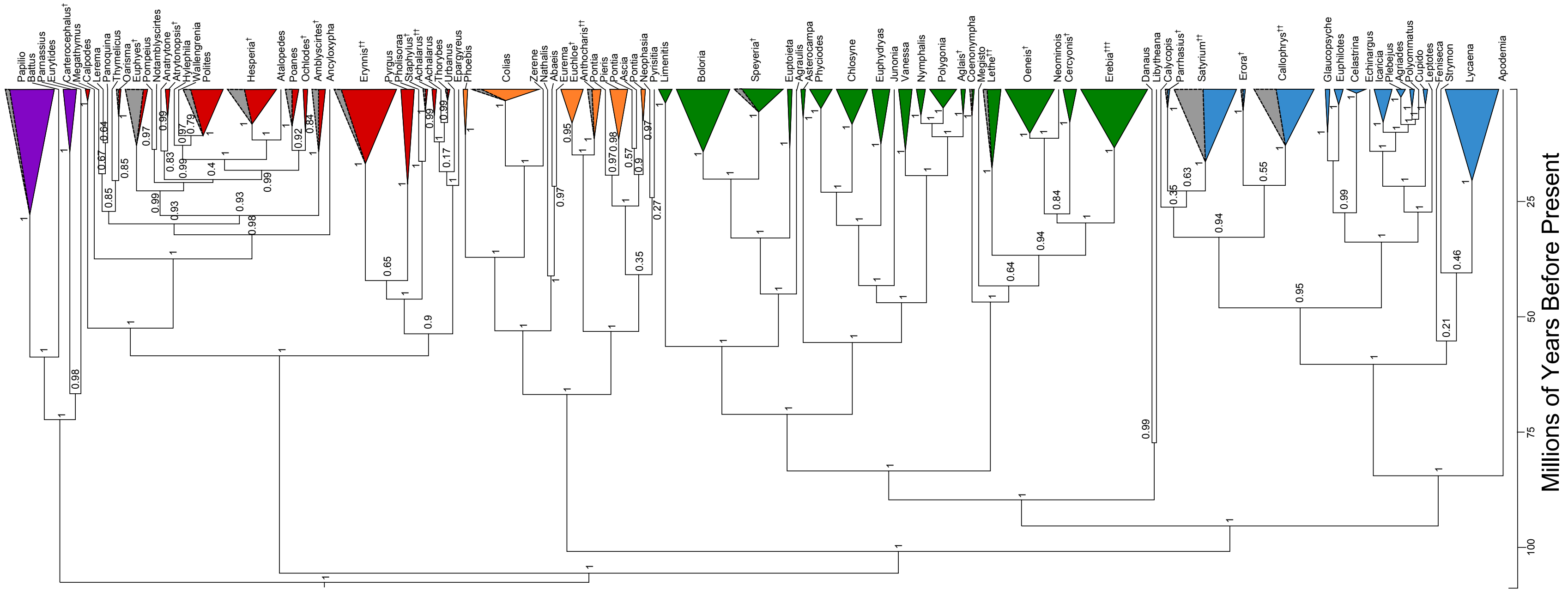

**Figure S1.** Complete, dated phylogeny of all 308 Canadian butterfly species and the 29 non-Canadian species used to help infer placement of data-deficient genera, collapsed to genus and coloured by family. The proportion of grey area per genus corresponds to number of data-deficient species per genus. Posterior probability values for each node are illustrated. † indicates a non-Canadian species that were used to place genera within the phylogeny; multiple † symbols indicate the number of non-Canadian species per genus. n=337 species.

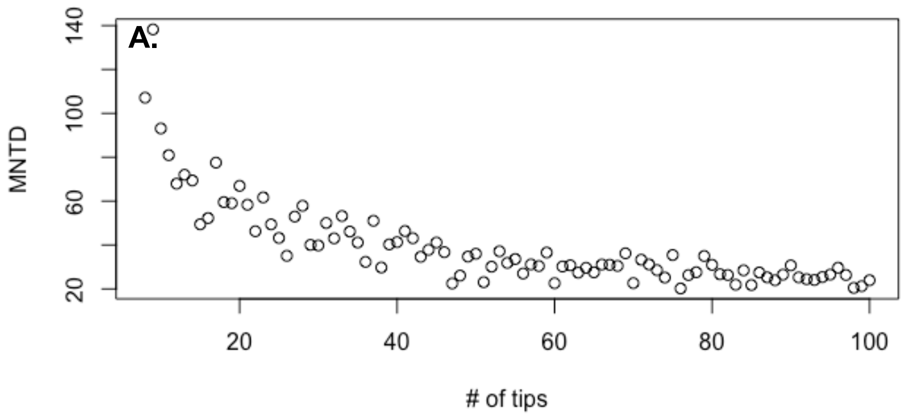

**Figure S2:** The relationship between Mean Nearest Taxon Distance (MNTD) and the number of tips in a community, drawn randomly from the full butterfly phylogeny.

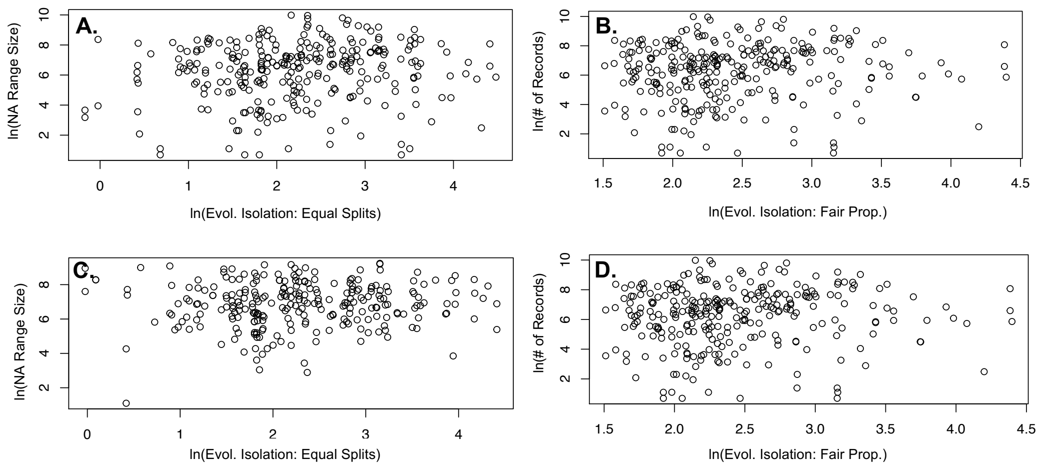

**Figure S3.** The relationship between a species evolutionary isolation score, measured as either equal splits (panels A, C) or fair proportions (B, D) and its rarity, measured as either the number of observations in Canada between 1900-2020 (A, B) or its' North American range size (C, D). ). Both isolation scores were generated using the MCC (Maximum Clade Credibility) version of the phylogeny. n = 289 species (A and B); 264 species (C and D).

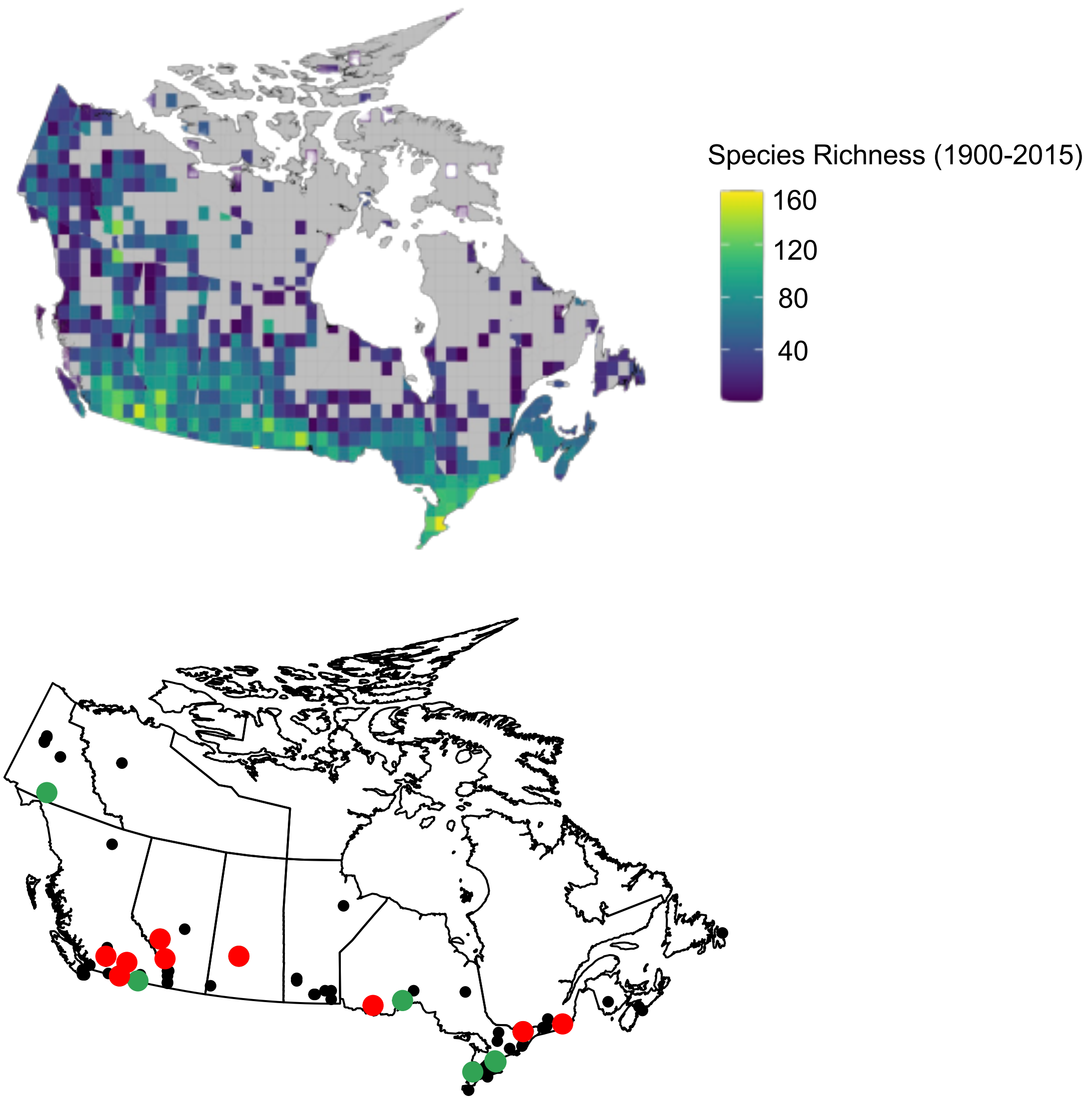

**Figure S4.** (A) Map of Chao-adjusted species richness across a 100 km x 100 km grid of Canada; and (B) Map of the top 10 grid cells experiencing the greatest increases and decreases in species richness across all quadrat/decade combinations. (A) shows species richness values across all grid cells, regardless of sampling intensity. (B) shows all grid cells that were adequately sampled and included in this study, where the 10 grid cells experiencing the greatest increase in species richness across all 10 x 10km quadrat/decade combinations are highlighted in green, and the 10 grid cells experiencing the greatest reduction in species richness are highlighted in red. n = 96 grid cells.

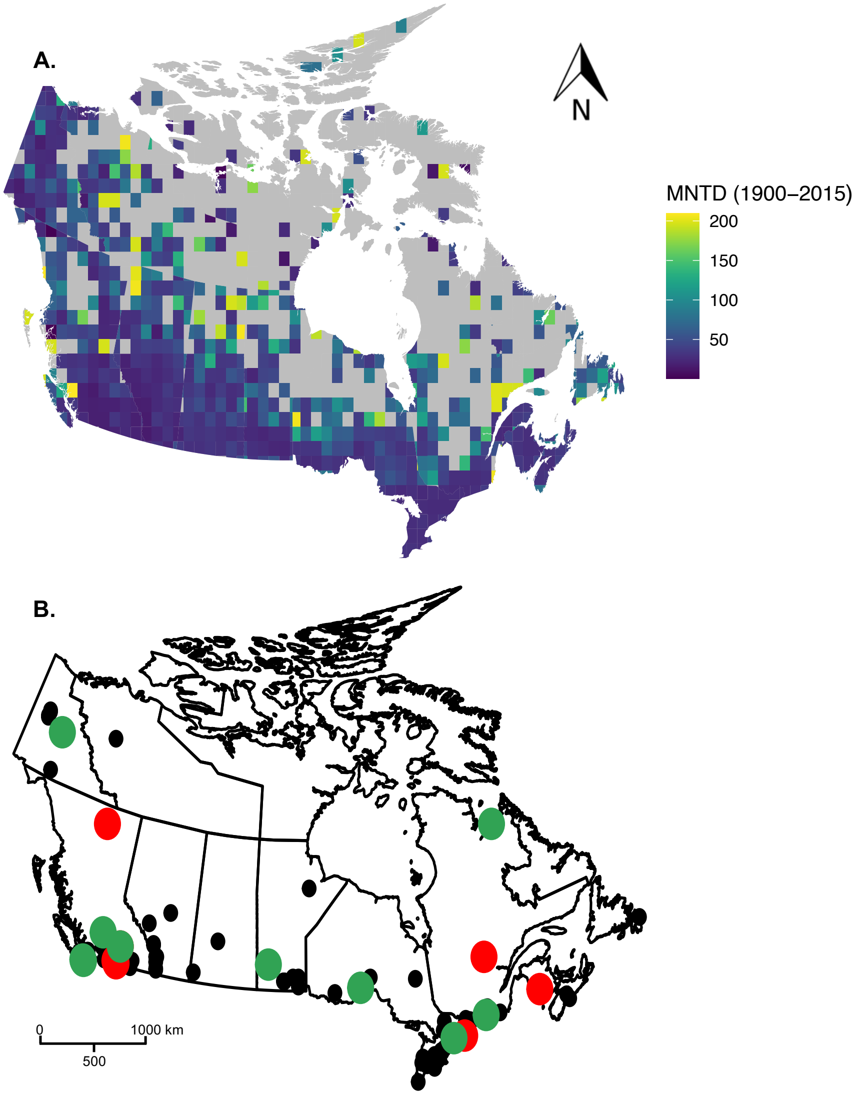

**Figure S5.** (A) Map of Mean Nearest Taxon Distance (MNTD) across a 100 km x 100 km grid of Canada; and (B) Map of the top 10 grid cells experiencing the greatest increases and decreases in MNTD across all quadrat/decade combinations
(A) shows MNTD values across all grid cells, regardless of sampling intensity. (B) shows all grid cells that were adequately sampled and included in this study, where the 10 grid cells experiencing the greatest increase in MNTD across all 10 x 10km quadrat/decade combinations are highlighted in green, and the 10 grid cells experiencing the greatest reduction in MNTD are highlighted in red. MNTD values were obtained by taking the average values across 1000 candidate tree topologies. n = 96 grid cells.

**Table S2.** Descriptive statistics of the biotic responses (change in species richness and change in Mean Nearest Taxon Distance) per decade.

| | Change in Species Richness | | | | Change in MNTD | | | |
|---|---|---|---|---|---|---|---|---|
| **Decade** | **Mean** | **SD** | **Median** | **n** | **Mean** | **SD** | **Median** | **n** |
| **1950s** | 6.99 | 25.62 | 9.05 | 37 | -4.39 | 62.94 | 2.68 | 39 |
| **1960s** | -0.14 | 27.18 | -0.05 | 45 | -2.57 | 55.95 | 1.78 | 53 |
| **1970s** | 15.06 | 25.04 | 15.15 | 68 | -23.49 | 48.92 | -12.13 | 79 |
| **1980s** | -0.53 | 25.27 | -0.25 | 83 | 8.19 | 32.11 | 2.66 | 87 |
| **1990s** | -10.03 | 29.66 | -9.80 | 69 | 10.88 | 36.47 | 8.24 | 74 |
| **2000s** | 7.06 | 35.18 | 1.67 | 56 | -3.94 | 46.43 | -1.57 | 63 |
| **2010s** | 11.81 | 32.87 | 12.48 | 61 | -12.97 | 48.99 | -5.62 | 69 |

SD = standard deviation, n = number of quadrats (out of a potential 96) with available data.

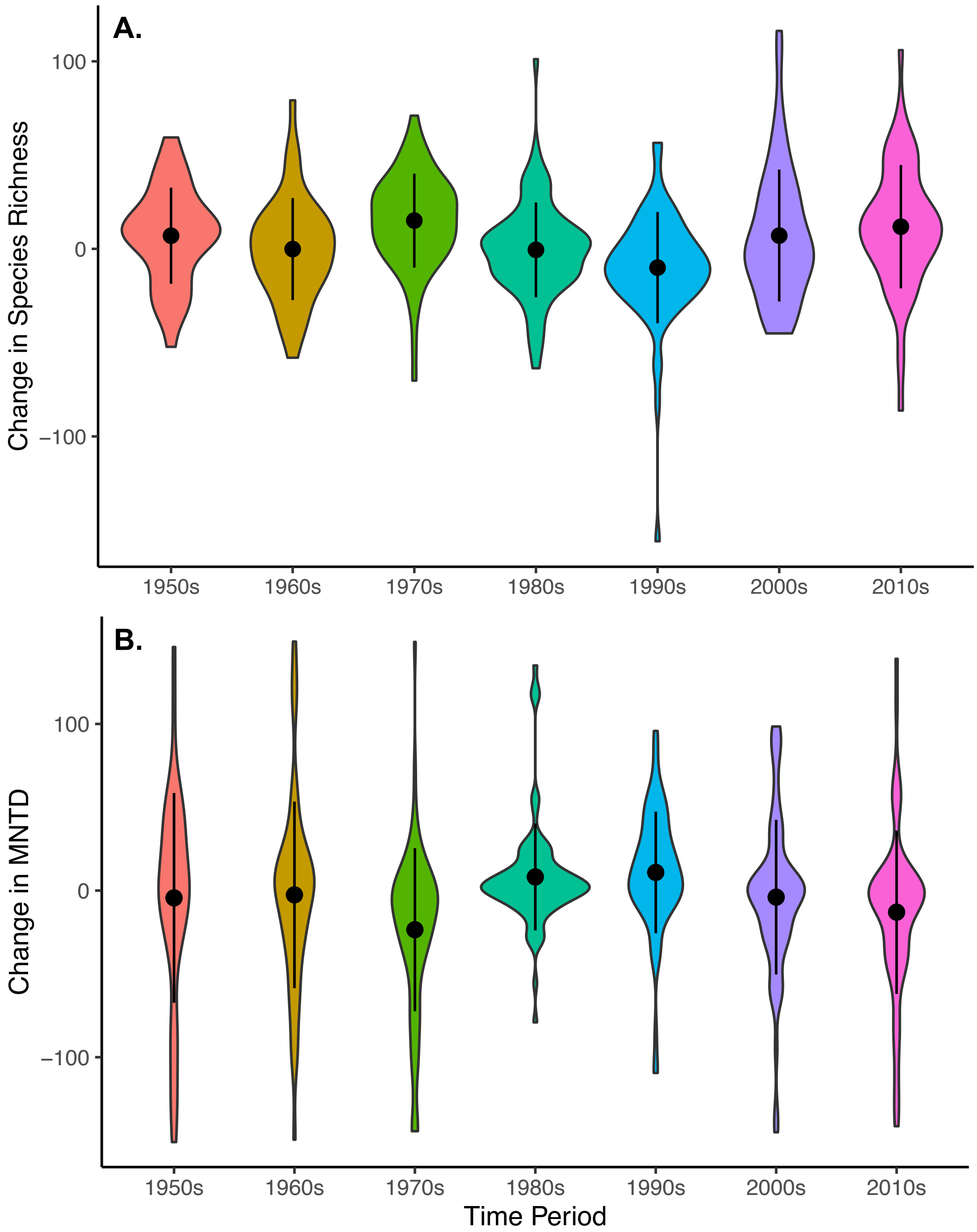

**Figure S6.** Violin plot of (A) the change in species richness and (B) the change in MNTD (averaged over 1000 candidate topologies) within a 10 km x 10 km grid cell, coloured by decade. See Table S2 for the number of quadrats per decade and per response.

### Supplementary Materials and Methods

### Part A: Choice of priors for Bayesian analysis of quadrat-level responses to land-use and climate change

Using each quadrat-level response per decade, I constructed two separate models: one to model change in species richness (hereafter "$\Delta$SR"), and one for change in MNTD (hereafter "$\Delta$MNTD").

Both response variables are numerical values between -155.92 to 116.22 (for $\Delta$SR) and -150.91 to 149.44 (for $\Delta$MNTD). As response variables were normally distributed, I used a Gaussian distribution with default priors to model their distributions.

The formula for each model was as follows:

$\Delta$SR ~ Hist. Primary to Crop TR + Cont. Primary to Crop. TR + Hist. Sec. to Crop. TR + Cont. Sec to Crop. TR + Hist. Crop to Sec. TR + Cont. Crop. To Sec. TR + Mean Temperature Change + (1|Quadrat ID)

$\Delta$MNTD ~ $\Delta$SR + Hist. Primary to Crop TR + Cont. Primary to Crop. TR + Hist. Sec. to Crop. TR + Cont. Sec to Crop. TR + Hist. Crop to Sec. TR + Cont. Crop. To Sec. TR + Mean Temperature Change + (1|Quadrat ID)

(where Hist. = Historic, Cont. = Contemporary, Crop. = Cropland, TR = Transition Rate)

I included 7 fixed effects and a random intercept for quadrat identity. All continuous predictors were standardized and centered at zero. I used weakly regularizing priors for all predictors, following recommendations in STAN (https://github.com/stan-dev/stan/wiki/Prior-Choice-Recommendations). For fixed effects, I used a diffuse Gaussian distribution with mean 0 and standard deviation of 3 for the slope. For the intercept, I again used a diffuse Gaussian prior with mean 0 and a standard deviation of 10. For the standard deviation associated with polygon identity, I used a Cauchy distribution that constraints SD to be positive, with a mean of 0 and SD of 1. All of these values are considered to be weakly informative, which is important because the idea is that a prior should "rule out unreasonable parameter values but is not so strong as to rule out values that might make sense" (https://github.com/stan-dev/stan/wiki/Prior-Choice-Recommendations).

I constructed 4 Markov chains of 4000 iterations each, including a warm-up phase of 1000 iterations. I used an adapt_delta value of 0.99 to reduce the number of divergent transitions. To verify convergence, I visually investigated the chains, as well as the $\hat{r}$ values. All $\hat{r}$ values were equal to 1, indicating model convergence.

### Part B: Habitat traits

I used 4 habitat-related traits in order to determine a species' sensitivity to land use change (Table S3). Although data was not available for all species, there was data for at least 205 species (out of 289) for some traits, and up to 261 species for others.

**Table S3.** The 4 traits used to determine habitat affinities, and their relevant trait states. Trait states were classified as categorical, but were ordered from one end of the trait spectrum to the other in order to measure phylogenetic signal in each trait. The final column shows the number of species in each category as well as the total number of species with data available.

| TRAIT | TRAIT VALUES | ORDERING | NUMBER OF SPECIES |
|---|---|---|---|
| **DISTURBANCE AFFINITY** | Disturbance-associated (strong) | 1 | 92 |
| | Disturbance-associated (weak) | 2 | 14 |
| | Disturbance association varies | 3 | 25 |
| | Seen near and away from disturbed habitat | 3 | 7 |
| | Disturbance-avoidant (weak) | 4 | 5 |
| | Disturbance-avoidant (strong) | 5 | 62 |
| | **TOTAL** | | **205** |
| **CANOPY COVER** | Closed canopy | 1 | 2 |
| | Mixed canopy (closed affinity) | 2 | 9 |
| | Canopy generalist | 3 | 61 |
| | Mixed canopy | 3 | 7 |
| | Mixed canopy (open affinity) | 4 | 118 |
| | Open canopy | 5 | 64 |
| | **TOTAL** | | **261** |
| **EDGE ASSOCIATION** | Edge-associated (strong) | 1 | 100 |
| | Edge-associated (weak) | 2 | 20 |
| | Edge association varies | 3 | 45 |
| | Seen near and away from edges | 3 | 11 |
| | No evidence for edge associations | 3 | 35 |
| | Edge-avoidant (weak) | 4 | 4 |

| | | | |
|---|---|---|---|
| | Edge-avoidant (strong) | 5 | 34 |
| | **TOTAL** | | **249** |
| **MOISTURE AFFINITY** | Mesic-associated (strong) | 1 | 109 |
| | Mesic-associated (weak) | 2 | 18 |
| | Both | 3 | 12 |
| | Moisture association varies | 3 | 28 |
| | No evidence for moisture association | 3 | 22 |
| | Xeric-associated (weak) | 4 | 8 |
| | Xeric-associated (strong) | 5 | 59 |
| | **TOTAL** | | **256** |

### Part C: Phylogeny Construction

#### i. Taxon Set and Taxonomic Data

Members of all six families of butterflies can be found in Canada (Papilionidae, Hesperiidae, Pieridae, Nymphalidae, Lycaenidae and Riodinidae). Species were selected for this tree after consulting the most recent taxonomy of North American butterflies (Pelham, 2014) and consulting the available literature and expert opinion as to which species are known to occur in Canada (Larrivée et al., 2014; Brock and Kaufman, 2003; Layberry et al, 1998).

A total of 308 species from Pelham (2014) were included in the final phylogeny (see Table S9 and S10 for a complete list of species). The taxonomy from Pelham (2014) was also used to assign taxon names and was the taxonomy used to place data-deficient species (see below for details on how data deficient species were added).

#### ii. Molecular Data Sampling

I used 8 genes to construct the phylogeny: one mitochondrial (COI) gene region and 7 protein-coding nuclear markers (*CAD, EF- 1α, GADPH, IDH, MDH, RpS5,* and *wingless*), totaling 7178 base pairs with gaps (see Table S4 for the length of each gene used and the frequency of each gene in the dataset). Although mtDNA data (specifically the COI gene) is more easily and widely available (Ratnasingham and Hebert, 2007) and has proven invaluable in the study of closely related taxa (including within Lepidoptera; Hajibabaei et al. 2007), because of the very low rate of recombination (Hebert et al. 2003), it also carries many potential pitfalls in phylogenetic analyses (see Rubinoff and Holland (2005) for a discussion of these). Thus, many authors advocate using it in concert with other independent genetic markers, such as nuclear DNA (Rubinoff and Holland 2005).

The large majority of my sequence data was obtained through GenBank (see Table S9 for ascension numbers). All of the sequence data used in the present study were downloaded on or before October 2018. Some nuclear data was provided by N. Wahlberg from previously published work (Heikkilä et al., 2012).

Available molecular data was unevenly distributed across genes and species. The vast majority of species had at least some data available for the COI (mitochondrial) gene, while molecular data for all nuclear genes was much more sparse (Table S4). Overall, 67.2% of the data matrix was missing nucleotides, with the MDH gene having the highest proportion of missing data (88.4%; Table S4). Additionally, some taxonomic groups were missing data for all genes for some members of their clade, while others had data for all members of their clades (Figure 4.2). For example, Papilionidae and Nymphalidae, both charismatic butterfly families had data for 94.4% and 95.1% of their taxonomic members, respectively, whereas Lycaenidae and Hesperiidae had data for 76.1% and 74.3% of their members (Table S5, Figure S1, Table S10).

Sequences were aligned in the multiple sequence alignment program MAFFT version 7 (Katoh and Standley, 2013), using the L-INS-i algorithm, which is considered to be the most accurate of the alignment methods (Katoh et al. 2005). I constructed individual gene trees in RAXML (Stamatakis, 2014) and examined the results for consistency across genes. Since there are many species that are completely lacking sequence data, I incorporated non-Canadian species (in the same genus) with sequence data to help infer the placement of these genera (Table S5). 294 species had data for at least one gene, 29 of which were non-Canadian species representing Canadian genera.

**Table S4.** Length (number of base pairs) of each gene used to construct the phylogeny, and frequency of each gene in the dataset.

| | COI | CAD | EF1-a | GADPH | IDH | MDH | RpS5 | wingless |
|---|---|---|---|---|---|---|---|---|
| **# of base pairs** | 1752 | 850 | 1282 | 721 | 716 | 732 | 616 | 502 |
| **% of base pairs missing** | 44.1% | 82.5% | 59.4% | 76.5% | 85.7% | 88.4% | 74.1% | 62.4% |
| **# of Canadian species** | 260 | 62 | 155 | 76 | 48 | 53 | 78 | 149 |
| **# of non-Canadian species** | 29 | 7 | 16 | 12 | 6 | 7 | 13 | 13 |

### iii. Initial Data Tree

A model of evolution was first determined using jModelTest (Darriba et al., 2012; Guidon and Gascuel, 2003) on individual gene regions. The model GTR + I + Γ consistently returned the lowest AIC score, and was therefore selected as the overall model of evolution.

All of my subsequent phylogeny construction was done using Bayesian inference in BEAST v1.10 (Suchard et al. 2018). In large analyses where there are a significant amount of missing genetic data, a starting tree (a tree from which an MCMC tree search is initialized) can greatly improve the speed and efficiency of the tree search. This starting tree is then modified each search iteration and evaluated to see if it is a better fit to the data.

The constant rate birth-death process is widely used as a prior distribution when modelling speciation and extinction. At any point in time during the process, each species has a constant probability of “dying” (going extinct), and a constant probability of giving “birth” (branching). However, the model is conditioned on observing *n* species at the present, and in order to estimate these instantaneous rates, it is assumed that all species which evolved under this model are sampled (Stadler, 2009). One of the challenges of building regional phylogenies is that they are, by definition, incomplete. In order to account for this, recent versions of BEAST include a parameter for “sampling probability” (the fraction of all living species within a clade that were sampled), which adjusts birth and death rates accordingly. I made use of that function here (see below).

Missing base pairs (bp) were coded as “?” and clock model and substitution models were left unlinked for individual gene regions and parameter values for each were estimated separately (i.e. a separate GTR + I + Γ model was specified for each partition). The clock prior was also unlinked between genes and set to uncorrelated relaxed clock to allow branch lengths to vary across lineages and genes (Drummond and Rambaut, 2007). The tree prior was set to a Birth-Death Incomplete Sampling process with default priors on growth and death rates. I used a beta distribution for the sampling probability so that it resulted in a bell-shaped curve that constrained 95% of the distribution to be between 0.008 and 0.025 with a median of 0.015. This would correspond to between 12,000 and 37,500 extant Papilionidae species; the median value of ~20,000 species is in line with current global diversity estimates (Espeland et al. 2018).

In order to satisfy downstream fossil calibrations, the starting tree must have nodes values (ages) that lie within the calibration constraints. As such, I used normal distributions to set an initial age value for the stem of Papilionidae (mean = 80 MYA, SD=0.1), the stem of Pieridae (mean = 80 MYA, SD=0.1), the stem of Nymphalidae (mean = 85 MYA, SD=0.1) and the stem of Lycaenidae (mean = 75 MYA, SD=0.1) using recent estimates of family ages (Espeland et al. 2018).

In order to make the tree search more efficient, I imposed a series of uncontroversial topological constraints following the most up-to-date knowledge of the phylogenetic relationships within butterflies (Espeland et al. 2018). Firstly, I constrained all families to be monophyletic. As for the order of branching of families, all except Papilionidae were constrained to form a monophyletic group; Pieridae, Nymphalidae, Lycaenidae and Riodinidae were constrained to be a monophyletic group; Nymphalidae, Lycaenidae and Riodinidae were constrained to be a monophyletic group; and Lycaenidae and Riodinidae were constrained to be a monophyletic group.

Using a series of hard topological constraints, 44 data-deficient species from across 20 genera were constrained to their relevant genus, but free to float around within that genus (see Table S10 for a list of data-deficient species).

In order to add data-deficient species to the phylogeny using these topological constraints, the genera that they belonged to had to be monophyletic. However, in early trial runs, some taxa were consistently placed outside of their genus. For these genera, I conducted Shimodaira-Hasegawa (SH) tests, which use the null hypothesis that all topologies are equally good representations of the data. Unconstrained data trees were compared to taxonomy-constrained trees for all genera that exhibited non-monophyly. If there was additional taxonomic information for the subgenus, subgenera were also tested. For all tests, there was no evidence of a significant difference in tree likelihoods between the taxonomy-constrained tree and the data tree. This indicated that the data could not reject taxonomic constraints, and so genus and subgenus monophyly constraints were enforced. A list of all the constraints on the topology can be found in Table S11.

In order to assess whether these taxonomic constraints had significant impact on node heights, I compared node ages for a topologically unconstrained tree to one that constrained those genera to be monophyletic. I extracted and compared the heights of all crown and stem nodes across all genera for the constrained and unconstrained trees. These taxonomic constraints did not seem to greatly influence my estimates of branching times however, as crown ages of genera on unconstrained trees were very similar to those on constrained trees (Figure S7a). However, stem ages of genera on constrained trees had some deviations from those obtained on unconstrained trees (Figure S7b). Most notably, the genus *Megathymus* has a stem age of 32.8 MYA on the unconstrained tree, and 16.8 MYA on the constrained topology. This genus has only one member in Canada (*Megathymus streckeri*), and this species has data for all 8 genes. Additionally, the node above this tip is well supported (posterior support of 1) on the final version of the phylogeny (Figure S1).

The search for the starting tree was run for 50,000 generations with parameters logged every 1000 generations. The maximum clade credibility tree was then extracted via TreeAnnotator v.1.10 (found within the BEAST package) and used as the initial tree for subsequent phylogenetic reconstructions.

Effective Sample Size (ESS; the number of effectively independent draws from the posterior distribution that the Markov chain is equivalent to) values were obtained in Tracer v.1.7.0 for three critical parameters: posterior (586), tree likelihood (1344) and root height (578). These values were above the best-practice threshold of 200, and thus indicated adequate posterior sampling.

**Table S5.** Taxonomic breakdown of species with and without genetic data, as well as the number of exotic species used in the phylogenetic inference of genus placement.

| Family | # of Can. species with data | # of Can. species missing data | Total Canadian species | # of exotic species used |
|---|---|---|---|---|
| **Hesperiidae** | 55 | 19 | 74 | 11 |
| **Lycaenidae** | 54 | 17 | 71 | 6 |
| **Nymphalidae** | 98 | 5 | 103 | 9 |
| **Papilionidae** | 17 | 1 | 18 | 0 |
| **Pieridae** | 39 | 2 | 41 | 3 |
| **Riodinidae** | 1 | 0 | 1 | 0 |
| **Total** | 265 | 44 | 308 | 29 |

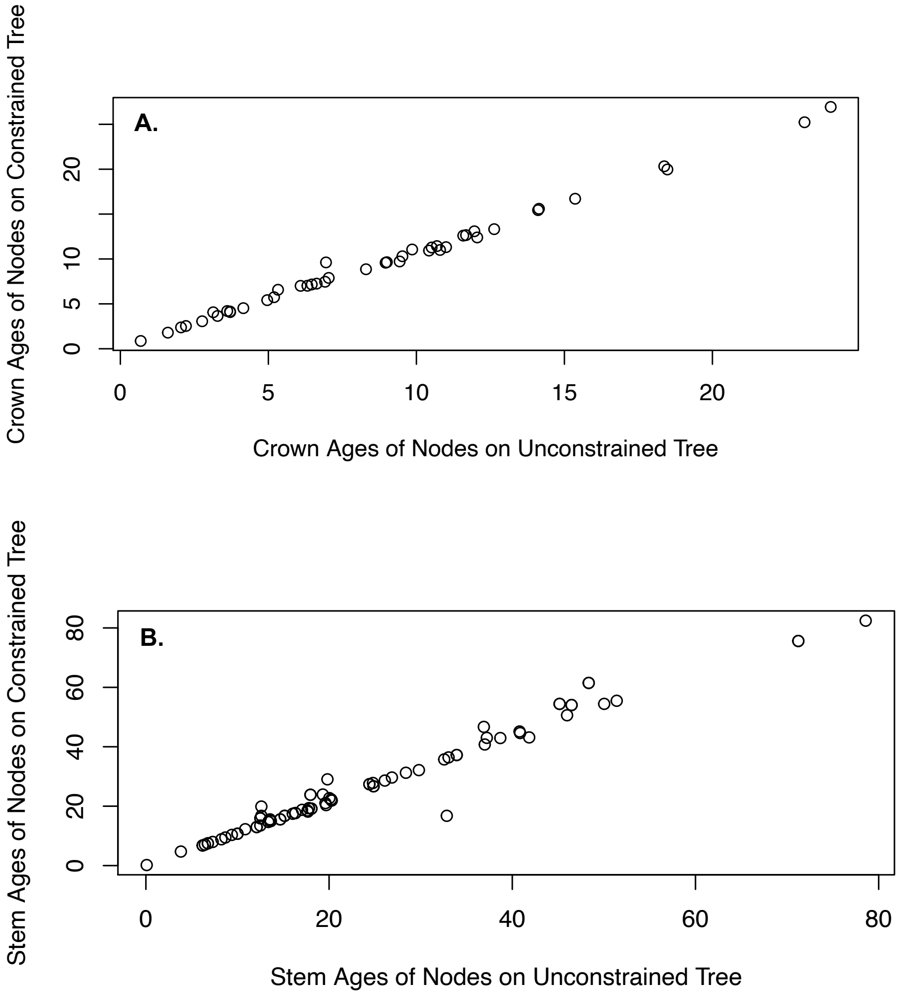

**Figure S7.** The relationship between the crown age (in millions of years) of all multitaxa genera (A) or stem age of all genera (B) on a phylogeny where genera were not constrained to be monophyletic vs. a phylogeny where 21 genera (see Table S11 for a list) were constrained to be monophyletic. (A) n = 44 nodes; (B) n = 85 nodes.

### iv. Dating the Phylogeny

After constructing the starting tree, I inferred the final phylogeny and times of divergence once again in BEAST 1.10. I used the same topological constraints, substitution model, clock model, tree prior (including birth rate, death rate and sampling probability) as in the starting tree. Additionally, I used a series of fossil calibrations in order to date the tree (see below for a full discussion of these). The search was conducted over 125 million generations with parameters logged every 10,000 generations and convergence statistics analysed using Tracer (Rambaut et al., 2018), with the first 25% of runs discarded as burn-in. Summary trees were obtained in TreeAnnotator v1.8 with the same 25% of runs discarded as burn-in, and a threshold on posterior probability was set at 50% support; nodes with support below this value become polytomies. In order to account for phylogenetic uncertainty in the placement of data-deficient species in downstream analyses, a distribution of 1000 possible tree topologies was generated by randomly sampling from the posterior distribution of tree topologies (after the burn-in).

Our estimates of crown ages for the 5 major families and the crown of butterflies were largely in line with what has been found in other studies (Table S8). Notably, our estimates tend to be younger than other, more global studies, which is to be expected as mine is a regional phylogeny and is missing representatives from many genera that aren’t found in Canada.

### v. Fossil Calibrations

The fossil record for Lepidoptera is notoriously poor (Sohn et al. 2015) but a few well-documented and well-established fossils were used to date the tree (see Table S6). Since we are not sampling entire families and thus cannot accurately estimate the age of the crown for most clades, all of my fossils were used to constrain the age of the stem of the group of interest instead of the crown (Forest, 2009). As recommended by several authors (i.e. Yang & Rannala, 2006), lognormal distributions were used for fossil priors; for specific values see Table S7. A hard minimum bound was set by the age of the fossil and a soft maximum bound was set by the origin of the angiosperms at 183 MYA (Bell et al. 2010), based on the assumption that all but the earliest Lepidoptera were constrained by the diversification of angiosperms (Condamine et al. 2012). A uniform prior was placed on the root of the tree, constraining it between 0 and 183 MYA (the origin of the angiosperms).

**Table S6.** List of relevant fossils used to help infer divergence times in the dated phylogeny.

| Fossil Name | Family | Date (hard minimum bound) | Reference | Estimated placement in the literature | Conservative placement in BEAST priors |
|---|---|---|---|---|---|
| ***Praepapilio*** | Papilionidae | 48 MYA | Condamine et al (2013) | Papilionidae family crown | Stem of Papilionidae |
| ***Nymphalites*** | Nymphalidae | 34 MYA | Emmel et al (1992) | Unclear; some similarities to modern-day *Marpesia*, *Anaea* and *Limenitis* genera | Stem of Nymphalidae |
| ***Stolopsyche libytheoides*** | Pieridae | 34 MYA | Braby et al. (2006); Heikkilä et al (2012) | Ancestor or sister genus to *Pieris* (within Pierini tribe) | Stem of Pieridae |
| ***Oligodonta florissantensis*** | Pieridae | 34 MYA | Braby et al. (2006); | Similar to the *Catasticta* and *Leodonta* genera | Stem of Pieridae |
| ***Thaites ruminianus*** | Papilionidae | 30 MYA | Condamine et al. (2012) | Sister to Luehdorfiini or Zerynthiini tribes (within Parnassinae subfamily) | Stem of Parnassinae |
| ***Coliates proserpina*** | Pieridae | 30 MYA | Braby et al. (2006) | Close relative of *Delias*, *Prioneris* and *Aporia* genera | Not used; older fossil used for stem of Pieridae |
| ***Vanessa americindica*** | Nymphalidae | 23 MYA | Emmel et al (1992) | Split between *Hypanartia* and *Vanessa* genera | Not used; older fossil used for stem of Nymphalidae |
| ***Miopieris talboti*** | Pieridae | 5.33 MYA | Braby et al. (2006) | Close relative of the *Pontia* genus | Not used; older fossil used for stem of Pieridae |
| ***Doritites bosniaskii*** | Papilionidae | 5.33 MYA | Condamine et al. (2012); Nazari et al. 2007 | Sister genus to *Archon* (Luehdorfiini tribe) | Not used; older fossil used for stem of Papilionidae |

Estimated fossil placements in the literature are compared to my relatively conservative priors enforced in BEAST.

**Table S7.** List of fossil constraints and priors specified in dated phylogeny.

| Fossil/Prior Name | Topological Constraint | Hard minimum bound | Soft maximum bound | Lognormal density mean | Lognormal density SD |
|---|---|---|---|---|---|
| ***Vanessa americindica*** | Stem of Nymphalidae | 34 MYA | 183 MYA | 108.5 (μ = 4.3108) | 0.42 |
| ***Praepapilio*** | Stem of Papilionidae | 48 MYA | 183 MYA | 115.5 (μ = 4.2121) | 0.42 |
| ***Thaites ruminianus*** | Stem of Parnassinae | 30 MYA | 183 MYA | 106.5 (μ = 4.337) | 0.42 |
| ***Stolopsyche libytheoides*** | Stem of Pieridae | 34 MYA | 183 MYA | 108.5 (μ = 4.3108) | 0.42 |

Parameters were specified so that the peak of the density is set by the mean (the difference between the hard minimum bound and the soft maximum bound). This standard deviation value splits the density to be 50:50 around the peak of the density, with 5% in the tail beyond the soft upper bound.

#### A. Papilionidae (*Praepapilio, Doritites bosniaskii*)

*Praepapilio colorado* and *P. gracilis* were first described by Durden and Rose (1978) in middle Eocene beds (which dates it between 38 and 48 million years ago (Cohen et al., 2013) and were considered to belong to the Papilionidae family. There are 3 subfamilies within Papilionidae: Baroniinae (usually considered the oldest of the subfamilies (Caterino et al., 2001)), Parnassinae and Papilionidae. *Praepapilio* has one synapomorphy of the Papilionidae (a basal spur) but lacks a single anal wing vein that is present in Papilionidae and thus shares a character state with Baroniinae and Parnassinae. Therefore, due to the ambiguity in characters, there is disagreement as to whether it should be basal to the whole Papilionidae family (de Jong, 2007) or only to the *Papilio* genus (Nazari et al., 2007). As a conservative estimate, I used the fossil to represent a minimum age of 48 MYA for the stem of Papilionidae.

*Doritites bosniaskii* is the sole member of its now-extinct genus, but morphological analyses place it sister to the modern-day *Archon* genus (Condamine et al. 2012; Nazari et al. 2007). As the *Archon* genus is not present in Canada, and since *Doritites bosniaskii* was dated to the late Miocene era (5.33-11.63 MYA), I decided not to use this fossil information in dating the phylogeny.

#### B. Parnassinae (*Thaites ruminianus*)

*Thaites ruminianus*, described by Scudder (1875), is another fossil from the Papilionidae family and is dated to the early Oligocene (28.4-33.9 MYA) (Condamine et al., 2012). It is an obvious member of the Parnassinae subfamily, however the precise position of *Thaites* within Parnassinae is unclear (Nazari et al., 2007) since it shares synapomorphies with 2 exclusive tribes within the subfamily. Therefore, it was used to provide a conservative minimum age of 30 Ma for the stem of Parnassinae. We enforced monophyly on the genus *Parnassius* in order to do so.

#### C. Nymphalidae (*Vanessa americindica, Nymphalites obscurum, Nymphalites scudderi*)

There are several fossils of the Nymphalidae family from the late Eocene (see Condamine et al., 2012 for examples). Scudder (1875) described *Nymphalites obscurum* and correctly placed it within Nymphalidae but its position within the family is less clear (Emmel et al., 1992) based on its unique combination of characters that are absent from any North American species. This specimen is dated to the late Eocene (38-34 MYA; Emmel et al., 1992). Similarly, *Nymphalites scudderi* bears some resemblance to modern-day Nymphalids but possesses some unique characters (such as the shape of the forewing) that make it difficult to place. This specimen is also dated to the late Eocene (Emmel et al. 1992). Therefore, I used the ages of these two fossils to constrain the minimum age of the Nymphalidae stem to 34 Ma.

*Vanessa amerindica*, a fossil only named in 1989, bears enough resemblance to the modern-day *Vanessa indica* (currently found in Asia) to be confidently placed within the *Vanessa* genus (Emmel et al. 1992). However, it is dated to the Oligocene (23-34 MYA), and so is younger than the *Nymphalites* fossils. As such, it was not used to constrain minimum ages in my analysis.

#### D. Pieridae (*Oligodonta florissantensis, Stolopsyche libytheoides, Coliates proserpina, Miopieris talboti*)

Similarly, there are a number of fossils from within Pieridae from the late Eocene period (see Heikkilä et al., 2012 and Braby et al., 2006), such as *Stolopsyche libytheoides* and *Oligodonta florissantensis*. *Stolopsyche libytheoides* is considered to be either the ancestor or sister taxon of the modern *Pieris* genus (a genus that is currently found in Canada). The placement of *Oligodonta florissantensis* is much less clear, and is likely related to the *Leodonta* and *Catasticta* genera (Braby et al. 2006), which are not found in Canada.

Other Pieridae fossils include *Coliates proserpina* and *Miopieris talboti* (Braby et al. 2006) but they are from the Lower Oligocene and Upper/Late Miocene epochs (30-33.5 and 5.33-11.63) MYA respectively), and so are not informative about the crown or stem age of Pieridae. Therefore, we used the *Stolopsyche libytheoides* and *Oligodonta florissantensis* fossils to constrain the minimum age of the Pieridae stem to 34 MYA.

**Table S8.** A comparison of estimates of crown ages (in millions of years) of 5 major butterfly families inferred here and in previous studies.

| Studies | Crown Ages of Butterfly Clades | | | | | |
|---|---|---|---|---|---|---|
| | Papilionoidea | Papilionidae | Hesperiidae | Pieridae | Nymphalidae | Lycaenidae |
| This study | 104.6 (81.5-131.4) | 70.1 (52.9-89.9) | 56.5 (42.8-71.3) | 64.8 (49.2-82.1) | 87.1 (67.11-109.5) | 58.3 (42.5-74.7) |
| Espeland et al. (2018) | 119 (91-143) | 84 (63-109) | 79 (60-99) | 87 (67-108) | 91 (71-112) | 78 (60-96) |
| Cong et al. (2017) | 133 (102-162) | N/A | 90 (33-113) | N/A | 87 (69-104) | N/A |
| Heikkila et al. (2012) | 110 (92-128) | 75 (62-88) | 65 (54-79) | 80 (67-97) | 87 (74-101) | 73 (58-84) |
| Wahlberg et al. (2009) | 104 (93-116) | 63 (52-76) | N/A | 73 (57-86) | 94 (84-104) | 75 (63-86) |

Credibility or confidence intervals, depending on dating method used, are shown in parentheses. Adapted from Espeland et al. (2018).

**Table S9:** List of species and their respective GenBank sequence accession numbers for the 8 genes used to construct the phylogeny.
* indicates sequences that were donated by Nikolas Walhberg
† indicates non-Canadian species that were used to place genera within the phylogeny and subsequently dropped from the final phylogeny.

| Species | CAD | COI | EF1-a | GADPH | IDH | MDH | RPS5 | wingless |
|---|---|---|---|---|---|---|---|---|
| *Abaeis nicippe* | - | GU089559 | - | - | - | - | - | - |
| †*Achalarus albociliatus* | - | KY019648.1 | - | - | - | - | - | - |
| *Achalarus lyciades* | - | NC_030602.1 :1241-2774 | - | - | - | - | - | - |
| †*Achalarus toxeus* | - | GU149308 | - | - | - | - | - | - |
| †*Aglais io* | *NW63-16 | KM592970.1: 1454-2984 | *NW63-16 | FJ639521 | - | *NW63-16 | FJ639576 | AF412766 |
| *Aglais milberti* | *NW77-14 | AY248787.1 | *NW77-14 | FJ639523 | *NW77-14 | *NW77-14 | FJ639578 | AY248828.1 |
| *Agraulis vanillae* | *NW152-16 | GQ864730.1 | DQ922873.1 | KP073355.1 | KP073632.1 | GQ865162.1 | GQ865383 | KF277395.1 |
| *Agriades glandon* | GQ128557.1 | EU330439.1 | EU326285.1 | - | - | - | - | GQ128839.1 |
| *Agriades optilete* | - | GQ129011 | - | - | - | - | - | - |
| †*Amblyscirtes exoteria* | KY045514.1 | KY019655.1 | EU364274.1 | KY027469.1 | KY027731.1 | KY027973.1 | KY028491. 1 | EU364072.1 |
| *Amblyscirtes vialis* | - | JF841299.1 | - | - | - | - | - | - |
| *Anatrytone logan* | KY045518.1 | KY019659.1 | EU364270.1 | KY027473.1 | KY027735.1 | KY027977.1 | KY028498. 1 | - |
| *Ancyloxypha numitor* | KY045521.1 | KY019662.1 | EU364258.1 | KY027476.1 | KY027738.1 | KY027980.1 | KY028498. 1 | - |
| †*Anthocharis belia* | KM046494.1 | KM046798.1 | AY870560.1 | KM046840.1 | KM046734.1 | KM046671.1 | KM046600. 1 | AY954604.1 |
| *Anthocharis sara* | - | KU874145.1 | - | - | - | - | - | - |
| †*Anthocharis scolymus* | - | GU372556.1 | GU372647.1 | - | - | - | - | - |

| Species | CAD | COI | EF1-a | GADPH | IDH | MDH | RPS5 | wingless |
|---|---|---|---|---|---|---|---|---|
| ***Apodemia mormo*** | - | NC_024571.1:1470-3005 | EU520324.1 | - | - | - | - | KT285983.1 |
| ***Ascia monuste*** | KM046498.1 | KM046802.1 | AY870582.1 | KM046845.1 | KM046738.1 | KM046678.1 | KM046605.1 | KM046565.1 |
| ***Asterocampa celtis*** | - | *NW120-2 | GQ864832 | - | - | - | - | *NW120-2 |
| ***Asterocampa clyton*** | - | AB501199.1:1-1544 | - | - | - | - | - | AF246556.1 |
| ***Atalopedes campestris*** | - | KY019674.1 | EU364267 | KY027486.1 | KY027748.1 | KY027992.1 | KY028507.1 | EU364067.1 |
| ***Atrytonopsis hianna*** | - | KT139715.1 | - | - | - | - | - | - |
| †***Atrytonopsis ovinia*** | - | DQ291894.1 | - | - | - | - | - | - |
| ***Battus philenor*** | - | AF170875.1:12-1542 | AF173415 | - | - | - | - | DQ351130.1 |
| ***Boloria alaskensis*** | | HQ161227 | HQ161299 | - | - | - | - | HQ161172 |
| ***Boloria alberta*** | - | HQ161243 | HQ161315 | - | - | - | - | HQ161186.1 |
| ***Boloria astarte*** | - | HQ161229.1 | HQ161301 | - | - | - | - | HQ161174.1 |
| ***Boloria bellona*** | - | HQ161290.1 | HQ161356 | - | - | - | - | AF246530 |
| ***Boloria chariclea*** | - | HQ161276.1 | HQ161333.1 | - | - | - | - | HQ161205.1 |
| ***Boloria epithore*** | - | AF170862.1:12-1542 | HQ161305 | - | - | - | - | HQ161177 |
| ***Boloria eunomia*** | - | HQ161271.1 | HQ161337.1 | - | - | - | - | HQ161202.1 |
| ***Boloria freija*** | - | HQ161273.1 | HQ161312 | - | - | - | - | HQ161203.1 |
| ***Boloria frigga*** | - | HQ161275 | HQ161341 | - | - | - | - | HQ161204 |
| ***Boloria improba*** | - | HQ161237.1 | HQ161293 | - | - | - | - | HQ161166 |
| ***Boloria natazhati*** | - | HQ161291 | HQ161357 | - | - | - | - | HQ161219 |
| ***Boloria polaris*** | - | HQ266652 | HQ266658.1 | - | - | - | - | HQ266646 |

| Species | CAD | COI | EF1-a | GADPH | IDH | MDH | RPS5 | wingless |
|---|---|---|---|---|---|---|---|---|
| ***Boloria selene*** | - | HQ161222.1 | HQ161294.1 | - | - | - | - | AY090134.1 |
| ***Callophrys affinis*** | - | HQ918886.1 | - | - | - | - | - | - |
| †***Callophrys chalybeitincta*** | - | JF810410.1 | - | - | - | - | - | - |
| ***Callophrys gryneus*** | - | KT133344.1 | - | - | - | - | - | - |
| ***Callophrys henrici*** | - | KP150273.1 | - | - | - | - | - | - |
| ***Callophrys irus*** | - | KP150261.1 | - | - | - | - | - | - |
| ***Callophrys johnsoni*** | - | KC473847.1 | - | - | - | - | JN000853.1 | - |
| ***Callophrys niphon*** | - | KP150297.1 | - | - | - | - | - | - |
| ***Callophrys polios*** | - | MF957135.1 | - | - | - | - | - | - |
| †***Callophrys rubi*** | - | HQ004146.1 | - | - | - | - | - | - |
| ***Callophrys spinetorum*** | - | KC473851.1 | - | - | - | - | JN000839.1 | - |
| ***Calpodes ethlius*** | KY045546.1 | KY019690.1 | EU364289 | KY027500.1 | - | KY028006.1 | - | JQ786715.1 |
| ***Calycopis cecrops*** | - | GU089704 | - | - | - | - | - | - |
| ***Carterocephalus palaemon*** | KY045552.1 | KY019697.1 | EU364183.1 | KY027507.1 | KY027763.1 | KY028014.1 | KY028529.1 | EU363990.1 |
| †***Carterocephalus silvicola*** | JN204936.1 | NC_024646.1:1484-3014 | JN204979.1 | - | JN205003.1 | JN205021.1 | JN205030.1 | JN204921.1 |
| ***Celastrina echo*** | - | GQ129018.1:1-1493 | GQ128707.1 | - | - | - | - | GQ128918.1 |
| ***Celastrina ladon*** | - | GU438796.1 | - | - | - | - | - | - |
| ***Celastrina lucia*** | - | HM415089.1 | - | - | - | - | - | - |
| ***Celastrina neglecta*** | EU141297.1 | EU141355.1 | EU136662.1 | EU141480.1 | EU141533.1 | EU141598.1 | - | EU141236.1 |
| ***Celastrina serotina*** | - | KY020378.1 | - | - | - | - | - | - |

| Species | CAD | COI | EF1-a | GADPH | IDH | MDH | RPS5 | wingless |
|---|---|---|---|---|---|---|---|---|
| †***Cercyonis meadii*** | - | GQ357250.1 | GQ357316.1 | GQ357511.1 | - | - | GQ357637.1 | GQ357383.1 |
| ***Cercyonis oetus*** | - | JF841058.1 | - | - | - | - | - | - |
| ***Cercyonis pegala*** | - | AY218239.1 | KT448650.1 | - | - | - | - | AY218277.1 |
| ***Cercyonis sthenele*** | - | KT126057.1 | - | - | - | - | - | - |
| ***Chlosyne acastus*** | - | AF187735.2 | *NW35-15 | - | - | - | - | AY788486.1 |
| ***Chlosyne damoetas*** | - | *NW122-13 | *NW122-13 | - | - | - | - | *NW122-13 |
| ***Chlosyne gorgone*** | - | AF187772.2 | *NW34-4 | - | - | - | - | AY788489.1 |
| ***Chlosyne harrisii*** | - | AF187773.2 | AY788729 | - | - | - | - | AY788490.1 |
| ***Chlosyne hoffmanni*** | - | KM042298.1 | KM042267.1 | - | - | - | - | KM042225.1 |
| ***Chlosyne nycteis*** | - | AF187788.2 | *NW34-5 | - | - | - | - | AY788493.1 |
| ***Chlosyne palla*** | - | AF187791.2 | AY788733 | - | - | - | - | AY788494.1 |
| ***Chlosyne whitneyi*** | - | KM042288.1 | KM042266.1 | - | - | - | - | KM042218.1 |
| ***Coenonympha tullia*** | - | AF170860.1:12-1542 | AF173399.2 | - | - | - | KF721248.1 | DQ351126.1 |
| ***Colias alexandra*** | - | KT144569.1 | - | - | - | - | - | - |
| ***Colias canadensis*** | - | EU583874.1 | - | - | - | - | - | - |
| ***Colias christina*** | - | JF841246.1 | - | - | - | - | - | - |
| ***Colias eurytheme*** | - | AF044024.1:11-1541 | AF173400.2 | - | - | - | - | AF537291.1 |
| ***Colias gigantea*** | - | JF841254.1 | - | - | - | - | - | - |
| ***Colias hecla*** | - | EU583868.1 | - | - | - | - | - | - |
| ***Colias interior*** | - | FJ851616 | FJ851622.1 | - | - | - | - | HM236309.1 |
| ***Colias meadii*** | - | EU583863.1 | FJ851628.1 | - | - | - | - | FJ851647.1 |
| ***Colias meadii elis*** | - | EU583862 | - | - | - | - | - | - |

| Species | CAD | COI | EF1-a | GADPH | IDH | MDH | RPS5 | wingless |
|---|---|---|---|---|---|---|---|---|
| ***Colias nastes*** | - | GU096735.1 | - | - | - | - | - | - |
| ***Colias occidentalis*** | - | KM540112.1 | - | - | - | - | - | - |
| ***Colias palaeno*** | GU828181.1 | GU828486.1 | GU829301.1 | GU829810.1 | GU830077.1 | GU830394.1 | GU830680.1 | GU829570.1 |
| ***Colias pelidne*** | - | FJ851617 | HM236306.1 | - | - | - | - | - |
| ***Colias philodice*** | KM046507.1 | EU583856.1 | DQ157890.1 | KM046854.1 | KM046747.1 | KM046684.1 | KM046613.1 | AF537292.1 |
| ***Colias tyche*** | - | HQ570192.1 | - | - | - | - | - | - |
| ***Cupido amyntula*** | - | JF841050.1 | - | - | - | - | - | - |
| ***Cupido comyntas*** | GQ128571.1 | GQ128954.1:1-1493 | GQ128643 | - | - | - | - | - |
| ***Danaus plexippus*** | *NW108-21 | DQ018954 | DQ018921 | EU141486 | EU141540 | EU141605 | EU141382 | DQ018891 |
| ***Echinargus isola*** | GQ128566.1 | DQ018947.1 | DQ018914.1 | - | - | - | - | DQ018885.1 |
| ***Epargyreus clarus*** | - | GU089840.1 | - | - | - | - | - | EU442878.1 |
| †***Erebia cassioides*** | - | KR138758.1 | - | KR139010.1 | - | - | KR138892.1 | KR139120.1 |
| ***Erebia disa*** | - | KR138841.1 | - | - | - | - | KR138971.1 | KR139165.1 |
| ***Erebia discoidalis*** | - | KR138807.1 | - | KR139048.1 | - | - | KR138937.1 | KR231860.1 |
| ***Erebia epipsodea*** | - | KT448682.1:1-1552 | KT448651.1 | KR139022.1 | - | - | KR138908.1 | KT448711.1 |
| †***Erebia euryale*** | KU577142.1 | KR138833.1 | - | KR139072.1 | KU577237.1 | KU577243.1 | KR138963.1 | KR231863.1 |
| ***Erebia fasciata*** | - | KU874992.1 | - | KR138999.1 | - | - | KR138880.1 | KR139107.1 |
| ***Erebia lafontainei*** | - | KR138795.1 | - | KR139038.1 | - | - | KR138923.1 | KR139152.1 |

| Species | CAD | COI | EF1-a | GADPH | IDH | MDH | RPS5 | wingless |
|---|---|---|---|---|---|---|---|---|
| ***†Erebia ligea*** | - | KR138753.1 | DQ338922.1 | KR138989.1 | - | - | KR138890.1 | KR139115.1 |
| ***Erebia mackinleyensis*** | - | KR138793.1 | - | KR139036.1 | - | - | KR138921.1 | KR139150.1 |
| ***Erebia magdalena*** | - | KR138777.1 | - | KR139023.1 | - | - | KR138909.1 | KR139135.1 |
| ***Erebia mancinus*** | - | KR138792.1 | - | KR139035.1 | - | - | KR138920.1 | KR139149.1 |
| ***Erebia occulta*** | - | KU874996.1 | - | - | - | - | - | - |
| ***Erebia pawloskii*** | - | KR138809.1 | - | KR139050.1 | - | - | KR138939.1 | KR139157.1 |
| ***Erebia rossii*** | - | KR138800.1 | - | KR138993.1 | - | - | KR138929.1 | KR139101.1 |
| ***Erebia vidleri*** | - | AB324843 | - | - | - | - | - | - |
| ***Erebia youngi*** | - | KR138791.1 | - | KR139033.1 | - | - | KR138918.1 | KR139147.1 |
| ***†Erora badeta*** | - | GU153153.1 | - | - | - | - | - | - |
| ***Erynnis afranius*** | KY045593.1 | KY019747.1 | EU364173.1 | KY027549.1 | - | KY028058.1 | KY028572.1 | EU363980.1 |
| ***Erynnis brizo*** | - | KT144025.1 | - | - | - | - | - | EU442873.1 |
| ***Erynnis funeralis*** | - | MF547081.1 | - | - | - | - | - | EU442866.1 |
| ***Erynnis horatius*** | KY045594.1 | KY019748.1 | EU364172.1 | KY027550.1 | - | KY028059.1 | KY028574.1 | EU363979.1 |
| ***Erynnis icelus*** | - | JF841281.1 | - | - | - | - | - | EU442869.1 |
| ***Erynnis juvenalis*** | - | KT622324.1 | - | - | - | - | - | - |
| ***Erynnis lucilius*** | - | KT128295.1 | - | - | - | - | - | - |
| ***Erynnis martialis*** | - | - | - | - | - | - | - | EU442868.1 |

| Species | CAD | COI | EF1-a | GADPH | IDH | MDH | RPS5 | wingless |
|---|---|---|---|---|---|---|---|---|
| **†*Erynnis montanus*** | - | NC_021427.1 :1508-3043 | - | - | - | - | - | - |
| ***Erynnis pacuvius*** | - | - | - | - | - | - | - | EU442864.1 |
| ***Erynnis persius*** | - | JF841280.1 | - | - | - | - | - | EU442853.1 |
| ***Erynnis propertius*** | - | FJ041310 | - | - | - | - | - | EU442835.1 |
| **†*Erynnis tristis*** | - | AF170858.1:12-1542 | AF173397.1 | - | - | - | - | EU442858.1 |
| ***Euchloe ausonides*** | KM046519.1 | KM046809.1 | AY870558.1 | KM046866.1 | KM046756.1 | KM046693.1 | KM046626.1 | KM046574.1 |
| ***Euchloe creusa*** | - | KU875022.1 | - | - | - | - | - | - |
| **†*Euchloe hyantis*** | - | FM196522 | - | - | - | - | - | - |
| ***Euchloe lotta*** | - | FR728201 | - | - | - | - | - | - |
| ***Euchloe naina*** | - | KU875023.1 | - | - | - | - | - | - |
| ***Euchloe olympia*** | - | KT127555.1 | - | - | - | - | - | - |
| ***Euphilotes ancilla*** | - | KC710407 | KC710444.1 | - | - | - | - | - |
| ***Euphilotes battoides*** | - | JF262047.1:1-1497 | AY675365.1 | - | - | - | - | - |
| ***Euphilotes enoptes*** | - | AY675410.1:1-1261 | AY675363.1 | - | - | - | - | GQ128921 |
| ***Euphydryas anicia*** | - | AF186926 | *NW11-7 | - | - | - | - | KJ906604 |
| ***Euphydryas chalcedona*** | - | AF187752.2 | AY788744 | - | - | - | - | KJ906604.1 |
| ***Euphydryas editha*** | - | AF187765.2 | AY788745.1 | - | - | - | - | AY788506.1 |
| ***Euphydryas gillettii*** | - | AF187771.2 | AY788746 | - | - | - | - | AY788507.1 |
| ***Euphydryas phaeton*** | GQ864647 | AF187797.2 | AY788747 | GQ864965 | - | GQ865208 | GQ865434 | AY788508.1 |
| **†*Euphyes peneia*** | - | GU155404 | - | - | - | - | - | - |

| Species | CAD | COI | EF1-a | GADPH | IDH | MDH | RPS5 | wingless |
|---|---|---|---|---|---|---|---|---|
| ***Euphyes vestris*** | KY045596.1 | KY019750.1 | EU364272.1 | KY027552.1 | KY027803.1 | KY028061.1 | KY028575.1 | - |
| ***Euptoieta claudia*** | GQ864650 | DQ922864.1 | DQ922896.1 | GQ864967 | GQ865095 | GQ865211 | GQ865437 | DQ922832 |
| ***Euptoieta hegesia*** | - | DQ922865.1 | DQ922897 | - | - | - | KM013207 | DQ922833 |
| ***Eurema mexicana*** | KM046520.1 | AY954568 | AY870563 | KM046868.1 | KM046758.1 | KM046695.1 | - | AY954598.1 |
| ***Eurytides marcellus*** | - | AF044022.1:17-1547 | AF044815 | - | - | - | - | DQ351128.1 |
| ***Feniseca tarquinius*** | KF787490.1 | KF787220.1 | - | KP215699.1 | - | - | - | - |
| ***Glaucopsyche lygdamus*** | KT286292.1 | FJ808849 | AY675364.1 | - | - | - | - | KT285981.1 |
| ***Glaucopsyche piasus*** | - | JF262056.1:1-1494 | JF271993 | - | - | - | - | HQ918001.1 |
| ***Hesperia assiniboia*** | - | HM415277.1 | - | - | - | - | - | - |
| ***Hesperia comma*** | - | HM860435.1 | - | - | - | - | - | AY700706.1 |
| ***Hesperia dacotae*** | - | JX679242 | - | - | - | - | - | - |
| †***Hesperia florinda*** | - | AB192493 | - | - | - | - | - | - |
| ***Hesperia juba*** | - | - | - | - | - | - | - | AY700754.1 |
| ***Hesperia leonardus*** | KY045611.1 | KY019768.1 | EU364265 | KY027568.1 | KY027815.1 | KY028076.1 | KY028586.1 | EU364065.1 |
| ***Hesperia nevada*** | - | - | - | - | - | - | - | AY700757.1 |
| ***Hesperia sassacus*** | - | KM553868.1 | - | - | - | - | - | - |
| ***Hylephila phyleus*** | KY045618.1 | AF170859.1:18-1548 | AF173398.1 | KY027575.1 | KY027822.1 | KY028084.1 | KY028591.1 | DQ351124.1 |
| ***Icaricia acmon*** | - | AF170864 | - | - | - | - | - | - |
| ***Icaricia icarioides*** | GQ128580.1 | GQ128963.1:1-1493 | GQ128652.1 | - | - | - | - | GQ128861.1 |
| ***Icaricia lupini*** | - | GQ128964 | - | - | - | - | - | - |

| Species | CAD | COI | EF1-a | GADPH | IDH | MDH | RPS5 | wingless |
|---|---|---|---|---|---|---|---|---|
| ***Icaricia saepiolus*** | GQ128583.1 | GQ128966.1:<br>1-1493 | GQ128655.1 | - | - | - | - | - |
| ***Icaricia shasta*** | GQ128584.1 | GQ128967.1:<br>1-1493 | GQ128656 | - | - | - | - | GQ128864.1 |
| ***Junonia coenia*** | KM013160 | NC_028207.1<br>:1490-2987 | EU053331.1 | KM013294 | - | KM013252 | - | KJ906605.1 |
| ***Leptotes marina*** | - | GQ129025.1:<br>1-1493 | GQ128714.1 | - | - | - | - | GQ128926 |
| ***Lerema accius*** | - | HQ583513.1 | - | - | - | - | - | - |
| ***Lethe anthedon*** | - | GQ357188.1 | GQ357257 | GQ357393 | - | - | GQ357522.<br>1 | GQ357322 |
| **†*Lethe diana*** | - | AB327283 | KM200194.1 | KM200250.1 | - | - | KM200275.<br>1 | KM200292.1 |
| ***Lethe eurydice*** | - | DQ338772.1 | DQ338914.1 | GQ357395.1 | - | - | GQ357524.<br>1 | DQ338621.1 |
| **†*Lethe minerva*** | EU141309.1 | DQ338768 | DQ338909.1 | EU141492.1 | EU141546.1 | EU1416x11.1 | EU141387.<br>1 | DQ338616.1 |
| ***Libytheana carinenta*** | GQ864673 | GQ864786.1 | GQ864880 | - | GQ865118 | - | GQ865462 | GQ864474 |
| ***Limenitis archippus*** | - | DQ205109.1:<br>1-1208 | DQ208217.1 | HQ291194 | HQ291220 | - | HQ291247 | GQ985317.1 |
| ***Limenitis arthemis*** | - | EU121990 | DQ208219 | HQ291196 | HQ291218 | - | HQ291249 | JQ786885.1 |
| ***Limenitis lorquini*** | - | DQ205106.1:<br>1-1208 | EF643326.1 | HQ291198 | HQ291225 | - | - | EU433944 |
| ***Limenitis weidemeyerii*** | - | DQ205101.1:<br>1-1208 | EF643347.1 | HQ291199 | HQ291228 | - | HQ291251.<br>1 | GQ985322.1 |
| ***Lycaena cupreus*** | - | FJ490471 | FJ490499.1 | - | - | - | - | - |
| ***Lycaena dione*** | - | FJ490480 | FJ490508.1 | - | - | - | - | - |
| ***Lycaena dorcas*** | - | FJ490485.1 | FJ490514.1 | - | - | - | - | - |

| Species | CAD | COI | EF1-a | GADPH | IDH | MDH | RPS5 | wingless |
|---|---|---|---|---|---|---|---|---|
| ***Lycaena dospassosi*** | - | HM436370.1 | FJ490515 | - | - | - | - | - |
| ***Lycaena editha*** | - | FJ490476 | FJ490504 | - | - | - | - | - |
| ***Lycaena epixanthe*** | - | HM415082.1 | - | - | - | - | - | - |
| ***Lycaena helloides*** | - | AY954562.1 | DQ018915.1 | - | - | - | - | AY954592.1 |
| ***Lycaena heteronea*** | - | EU330432 | EU326289.1 | - | - | - | - | - |
| ***Lycaena hyllus*** | - | FJ490479 | - | - | - | - | - | - |
| ***Lycaena mariposa*** | - | FJ490487 | FJ490516.1 | - | - | - | - | - |
| ***Lycaena nivalis*** | - | EU330433 | EU326288.1 | - | - | - | - | - |
| ***Lycaena phlaeas*** | - | NC_023087.1 :1463-2969 | FJ490517 | AB696721.1 | - | - | - | - |
| ***Lycaena rubidus*** | - | EU330442.1 | FJ490495 | - | - | - | - | - |
| ***Megathymus streckeri*** | KY045632.1 | EU364504.1 | EU364299 | KY027589.1 | KY027836.1 | KY028099.1 | KY028606. 1 | EU364094 |
| ***Megisto cymela*** | *CP21-04 | GQ864789.1 | AY509084.1 | GQ864996.1 | *CP21-04 | *CP21-04 | GQ357569. 1 | GQ357341.1 |
| ***Nathalis iole*** | KM046542.1 | AY954569.1 | AY870562.1 | - | KM046775.1 | KM046714.1 | KM046648. 1 | AY954599.1 |
| ***Neominois ridingsii*** | - | DQ338870 | DQ339026.1 | - | - | - | - | DQ338735 |
| ***Neophasia menapia*** | - | KM046824.1 | AY870536.1 | KM046890.1 | - | - | KM046649. 1 | DQ082814.1 |
| ***Notamblyscirtes simius*** | KY045515.1 | KY019656.1 | EU364275.1 | KY027470.1 | KY027732.1 | KY027974.1 | KY028492. 1 | EU364073.1 |
| ***Nymphalis antiopa*** | - | AY218246.1 | *NW70-2 | FJ639524 | - | - | FJ639579 | AY218284.1 |
| ***Nymphalis californica*** | *NW74-14 | AY248789.1 | *NW74-14 | FJ639525 | *NW74-14 | *NW74-14 | FJ639580 | AY248830.1 |
| ***Nymphalis l-album*** | *NW78-1 | AY248791 | *NW78-1 | FJ639526 | *NW78-1 | *NW78-1 | FJ639581 | AY248832 |

| Species | CAD | COI | EF1-a | GADPH | IDH | MDH | RPS5 | wingless |
|---|---|---|---|---|---|---|---|---|
| ***Oarisma garita*** | KY045659.1 | KY019824.1 | EU364259.1 | - | - | KY028124.1 | KY028638.1 | EU364059.1 |
| ***†Ochlodes ochracea*** | - | AB192492 | KM669655.1 | - | - | - | - | - |
| ***Ochlodes sylvanoides*** | KY045660.1 | KY019825.1 | DQ018902.1 | KY027616.1 | KY027861.1 | KY028125.1 | KY028639.1 | DQ018872.1 |
| ***Oeneis alberta*** | - | FJ808851 | - | - | - | - | - | - |
| ***Oeneis alpina*** | - | LC155480.1 | LC155660.1 | - | - | - | - | - |
| ***Oeneis bore*** | - | LC155481.1 | LC155661.1 | KP888702.1 | - | - | KP888737.1 | KP888775.1 |
| ***Oeneis chryxus*** | - | *NW79-1 | EU326283.1 | KP888713.1 | - | - | KP888748.1 | KP888788.1 |
| ***Oeneis jutta*** | - | DQ018958.1 | DQ018925.1 | GQ357506.1 | - | - | GQ357632.1 | DQ018896.1 |
| ***Oeneis macounii*** | - | *NW79-5 | - | - | - | - | - | - |
| ***Oeneis melissa*** | - | KP888679.1 | LC155637.1 | KP888719.1 | - | - | - | KP888797.1 |
| ***Oeneis nevadensis*** | - | *NW79-9 | - | - | - | - | - | - |
| ***†Oeneis norna*** | - | LC155450.1 | LC155630.1 | KP888725.1 | - | - | KP888757.1 | KP888802.1 |
| *Oeneis polixenes* | - | LC155482.1 | LC155662.1 | - | - | - | - | - |
| ***Oeneis uhleri*** | - | FJ808863 | LC155665.1 | - | - | - | KP888766.1 | KP888811.1 |
| ***Panoquina ocola*** | KY045669.1 | KY019837.1 | EU364290 | KY027625.1 | KY027868.1 | KY028134.1 | KY028647.1 | - |
| ***Papilio brevicauda*** | - | HM416180.1 | - | - | - | - | - | - |
| ***Papilio canadensis*** | - | AF044014.1:11-1541 | AF044816.2 | - | - | - | - | AY569125.1 |
| ***Papilio cresphontes*** | - | AF044004.1:11-1541 | AF044832.1 | - | - | - | - | - |

| Species | CAD | COI | EF1-a | GADPH | IDH | MDH | RPS5 | wingless |
|---|---|---|---|---|---|---|---|---|
| ***Papilio glaucus*** | EU141334.1 | NC_027252.1:1202-2732 | AF044826.1 | EU141512.1 | EU141571.1 | EU141635.1 | EU141412.1 | AF233563.2 |
| ***Papilio indra*** | - | AF044011.1:11-1541 | AF044824 | - | - | - | - | - |
| ***Papilio machaon*** | - | AF044006.1:11-1541 | - | - | - | - | - | AY569124.1 |
| ***Papilio multicaudata*** | - | EF126475 | - | - | - | - | - | - |
| ***Papilio polyxenes*** | - | AF044010.1:11-1541 | AF044823.2 | - | - | - | - | - |
| ***Papilio rutulus*** | - | AF044015.1:11-1541 | AY954620.1 | - | - | - | - | - |
| ***Papilio troilus*** | - | AF044017.1:11-1541 | AF423810.1 | - | - | - | - | - |
| ***Papilio zelicaon*** | - | AF044008.1:11-1541 | AF044827.1 | - | - | - | - | - |
| ***Parnassius clodius*** | - | AF170871.1:12-1542 | AF173411.2 | - | - | - | - | DQ351134.1 |
| ***Parnassius eversmanni*** | - | EU093022.1 | EF485079 | - | - | - | - | - |
| ***Parnassius phobeus*** | - | JN204959 | - | - | - | - | - | - |
| ***Parnassius smintheus*** | - | LT999983.1:1448-2978 | EF485052.1 | - | - | - | - | AY569045.1 |
| †***Parrhasius polibetes*** | - | JQ550019 | - | - | - | - | - | - |
| ***Phoebis philea*** | - | EU583850.1 | - | - | - | - | - | - |
| ***Phoebis sennae*** | KM046547.1 | KM046828.1 | AY870571 | KM046896.1 | KM046780.1 | KM046719.1 | KM046655.1 | - |
| ***Pholisora catullus*** | KY045683.1 | EU364386.1 | KP895765.1 | KY027637.1 | KY027881.1 | KY028149.1 | KY028664.1 | EU363988.1 |
| ***Phyciodes batesii*** | - | AF187747.2 | EF494005 | - | - | - | - | EF493898.1 |

| Species | CAD | COI | EF1-a | GADPH | IDH | MDH | RPS5 | wingless |
|---|---|---|---|---|---|---|---|---|
| ***Phyciodes cocyta*** | GQ864697 | AF187755.2 | *NW11-4 | - | - | GQ865253.1 | GQ865486 | AY090158.1 |
| ***Phyciodes mylitta*** | - | AF187785.2 | AY788791 | - | - | - | - | AY788551.1 |
| ***Phyciodes pallida*** | - | AY156636.1 | AY788794 | - | - | - | - | EF493904.1 |
| ***Phyciodes pulchella*** | - | AF187783.2 | AY788797 | - | - | - | - | EF493906.1 |
| ***Phyciodes tharos*** | - | AF187807.2 | AY788798 | - | - | - | - | EF493903.1 |
| ***Pieris angelika*** | - | KT129178.1 | - | - | - | - | - | - |
| ***Pieris marginalis*** | - | JF841232.1 | - | - | - | - | - | - |
| ***Pieris oleracea*** | - | JF841233.1 | - | - | - | - | - | - |
| ***Pieris rapae*** | - | JN204969.1 | AY870550.1 | JN204991.1 | JN205008.1 | JN205026.1 | JN205036.1 | AY954611.1 |
| ***Pieris virginiensis*** | - | KT133944.1 | - | - | - | - | - | - |
| ***Plebejus anna*** | GQ128589.1 | GQ128972.1: 1-1493 | GQ128661.1 | - | - | - | - | GQ128869.1 |
| ***Plebejus idas*** | - | GQ128973 | - | - | - | - | - | - |
| ***Plebejus melissa*** | GQ128593.1 | GQ128975.1: 1-1493 | GQ128664.1 | - | - | - | - | GQ128873.1 |
| ***Poanes hobomok*** | - | JF841270.1 | - | - | - | - | - | - |
| ***Poanes zabulon*** | - | GU090121.1 | - | - | - | - | - | - |
| ***Polites draco*** | - | KT125949.1 | - | - | - | - | - | - |
| ***Polites mystic*** | - | JF841290.1 | - | - | - | - | - | - |
| ***Polites origenes*** | - | KT138122.1 | - | - | - | - | - | - |
| ***Polites peckius*** | - | HM415292.1 | - | - | - | - | - | - |
| ***Polites rhesus*** | - | KT125275.1 | - | - | - | - | - | - |
| ***Polites sabuleti*** | - | - | - | - | - | - | - | AY700751.1 |

| Species | CAD | COI | EF1-a | GADPH | IDH | MDH | RPS5 | wingless |
|---|---|---|---|---|---|---|---|---|
| ***Polites themistocles*** | KY045687.1 | EU364471.1 | EU364266.1 | KY027641.1 | KY027885.1 | KY028153.1 | KY028668.1 | EU364066.1 |
| ***Polites vibex*** | - | MF545442.1 | - | - | - | - | - | - |
| ***Polygonia comma*** | *NW65-6 | AY248794.1 | *NW65-6 | FJ639512 | *NW65-6 | *NW65-6 | FJ639567 | FJ639370 |
| ***Polygonia faunus*** | *NW74-12 | AY248798.1 | FJ650367.1 | FJ650382.1 | *NW74-12 | *NW74-12 | FJ639571 | AY248837.1 |
| ***Polygonia gracilis*** | *EW22-8 | AY248797.1 | FJ650370.1 | FJ639503 | *EW22-8 | *EW22-8 | FJ639558 | AY248836.1 |
| ***Polygonia interrogationis*** | *NW77-12 | AY248793,1 | *NW77-12 | FJ639519.1 | *NW77-12 | *NW77-12 | FJ639574 | AY248834.1 |
| ***Polygonia oreas*** | - | FJ639421.1 | FJ639477.1 | FJ650388.1 | - | - | FJ639570 | AY788561 |
| ***Polygonia progne*** | *EW21-4 | AY248795.1 | FJ639480.1 | FJ639502 | - | *EW21-4 | FJ639557 | FJ639375.1 |
| ***Polygonia satyrus*** | *NW74-9 | AY248796,1 | FJ650378.1 | FJ650392.1 | *NW74-9 | *NW74-9 | FJ639573 | AY248835.1 |
| ***Polyommatus icarus*** | GQ128611.1 | AY496817.1:1-1257 | AY496846.1 | - | - | - | - | GQ128891.1 |
| ***Pompeius verna*** | - | GU090137.1 | - | - | - | - | - | - |
| ***Pontia beckerii*** | - | EU583849.1 | - | - | - | - | - | - |
| ***Pontia occidentalis*** | - | JF841231.1 | - | - | - | - | - | - |
| ***Pontia protodice*** | - | KT133037.1 | - | - | - | - | - | - |
| ***Pontia sisymbrii*** | - | AF044890 | - | - | - | - | - | - |
| ***Pyrgus centaureae*** | - | HM430256.1 | - | - | - | - | - | - |
| ***Pyrgus communis*** | KY045701.1 | AF170857.1:16-1546 | AF173396.1 | KY027653.1 | KY027897.1 | KY028166.1 | KY028680.1 | AY569043.1 |
| ***Pyrgus ruralis*** | KY045699.1 | EU364382.1 | EU364177.1 | KY027654.1 | - | KY028167.1 | - | EU363984.1 |
| ***Pyrgus scriptura*** | KY045700.1 | KY019874.1 | EU364178.1 | KY027655.1 | - | KY028168.1 | KY028681.1 | EU363985.1 |
| ***Pyrisitia lisa*** | - | HQ583574.1 | - | - | - | - | - | - |
| ***Satyrium behrii*** | - | EU330438 | EU326284.1 | - | - | - | - | - |

| Species | CAD | COI | EF1-a | GADPH | IDH | MDH | RPS5 | wingless |
|---|---|---|---|---|---|---|---|---|
| ***Satyrium calanus*** | - | KT145203.1 | - | - | - | - | - | - |
| ***Satyrium lipaprops*** | - | FJ808939 | - | - | - | - | - | - |
| †***Satyrium pruni*** | - | GU372540.1 | KM211588.1 | - | - | - | - | - |
| ***Satyrium sylvinus*** | - | KM554247.1 | - | - | - | - | - | - |
| ***Satyrium titus*** | - | FJ808947 | - | - | - | - | - | - |
| †***Satyrium w-album*** | - | GU372568 | KM211586.1 | - | - | - | - | - |
| ***Speyeria aphrodite*** | - | JF841091.1 | - | - | - | - | - | - |
| ***Speyeria atlantis*** | - | HM414906.1 | - | - | - | - | - | - |
| ***Speyeria callippe*** | - | KT135463.1 | - | - | - | - | - | - |
| ***Speyeria cybele*** | - | DQ922863.1 | DQ922895 | - | - | - | - | DQ922831 |
| †***Speyeria diana*** | - | KY773334.1 | KY773378.1 | KY773438.1 | - | - | KY773542.1 | KY773482.1 |
| ***Speyeria edwardsii*** | - | KY773311.1 | KY773355.1 | KY773409.1 | - | - | KY773513.1 | KY773459.1 |
| ***Speyeria hydaspe*** | - | KT127948.1 | - | - | - | - | - | - |
| ***Speyeria idalia*** | - | KY773332.1 | KY773376.1 | KY773436.1 | - | - | KY773540.1 | KY773480.1 |
| ***Speyeria mormonia*** | - | EU330436 | EU326287.1 | - | - | - | - | - |
| ***Speyeria zerene*** | - | *NW138-14 | - | - | - | - | - | - |
| †***Staphylus ceos*** | KY045723.1 | KY045723.1 | EU364151 | KY027680.1 | KY027922.1 | KY028193.1 | KY028703.1 | EU363958 |
| ***Strymon melinus*** | - | GU090202.1 | - | - | - | - | - | - |
| ***Thorybes pylades*** | KY045738.1 | EU364331 | EU364126.1 | KY027696.1 | KY027938.1 | KY028210.1 | KY028716.1 | EU442877.1 |
| ***Thymelicus lineola*** | JN204935.1 | JN204962.1 | JN204978.1 | JN204987.1 | JN205002.1 | JN205020.1 | JN205029.1 | JN204920.1 |

| Species | CAD | COI | EF1-a | GADPH | IDH | MDH | RPS5 | wingless |
|---|---|---|---|---|---|---|---|---|
| ***Urbanus proteus*** | - | DQ293840.1 | - | - | - | - | - | - |
| ***Vanessa annabella*** | HQ734872.1 | HQ734905.1 | KJ648965.1 | HQ734970.1 | HQ734992.1 | HQ735014.1 | HQ735046.1 | KJ649141.1 |
| ***Vanessa atalanta*** | GQ864722 | HQ734886.1 | *NW63-21 | GQ865045 | GQ865155 | GQ865275 | HQ735029.1 | KJ649120.1 |
| ***Vanessa cardui*** | HQ734873 | EF683677.1 | HQ734947.1 | HQ734971 | HQ734993 | HQ735015 | HQ735047 | KX824730.1 |
| ***Vanessa virginiensis*** | KT286296.1 | AY248783.1 | *NW77-16 | HQ734973 | HQ734995 | KJ649034.1 | HQ735033.1 | AY248827.1 |
| ***Wallengrenia egeremet*** | - | GU090235.1 | - | - | - | - | - | - |
| ***Zerene cesonia*** | - | KM046838.1 | AY870567.1 | - | KM046796.1 | KM046731.1 | KM046668.1 | KM046597.1 |

**Table S10:** Species missing genetic data, and added using taxonomic constraints

| Species Missing Genetic Data |
|---|
| *Amblyscirtes hegon* |
| *Amblyscirtes oslari* |
| *Anthocharis stella* |
| *Callophrys augustinus* |
| *Callophrys eryphon* |
| *Callophrys fotis* |
| *Callophrys lanoraieensis* |
| *Callophrys mossii* |
| *Callophrys nelsoni* |
| *Callophrys sheridanii* |
| *Coenonympha nipisiquit* |
| *Colias johanseni* |
| *Erora laeta* |
| *Erynnis baptisiae* |
| *Erynnis zarucco* |
| *Euphyes bimacula* |
| *Euphyes conspicua* |
| *Euphyes dion* |
| *Euphyes dukesi* |
| *Hesperia colorado* |
| *Hesperia ottoe* |
| *Hesperia pahaska* |
| *Hesperia uncas* |
| *Lethe appalachia* |
| *Oarisma powesheik* |
| *Oeneis philipi* |
| *Papilio eurymedon* |
| *Parrhasius m-album* |
| *Poanes massasoit* |
| *Poanes viator* |
| *Polites baracoa* |
| *Polites sonora* |
| *Satyrium acadica* |
| *Satyrium californica* |
| *Satyrium caryaevorus* |
| *Satyrium edwardsii* |

| **Species Missing Genetic Data** |
|---|
| *Satyrium fuliginosa* |
| *Satyrium saepium* |
| *Satyrium semiluna* |
| *Speyeria coronis* |
| *Speyeria hesperis* |
| *Staphylus hayhurstii* |
| *Thorybes bathyllus* |

**Table S11.** List of groups that were constrained in the construction of the phylogeny.
† indicates non-Canadian species that were used to place genera within the phylogeny and subsequently dropped from the final phylogeny
*indicates a species missing genetic data

| Hesperiidae (Family) | Nymphalidae (Family) | Lycaenidae (Family) |
|---|---|---|
| †*Achalarus albociliatus* | †*Aglais io* | *Agriades glandon* |
| *Achalarus lyciades* | *Aglais milberti* | *Agriades optilete* |
| †*Achalarus toxeus* | *Agraulis vanillae* | *Callophrys affinis* |
| †*Amblyscirtes exoteria* | *Asterocampa celtis* | **Callophrys augustinus* |
| **Amblyscirtes hegon* | *Asterocampa clyton* | †*Callophrys chalybeitincta* |
| **Amblyscirtes oslari* | *Boloria alaskensis* | **Callophrys eryphon* |
| *Amblyscirtes vialis* | *Boloria alberta* | **Callophrys fotis* |
| *Anatrytone logan* | *Boloria astarte* | *Callophrys gryneus* |
| *Ancyloxypha numitor* | *Boloria bellona* | *Callophrys henrici* |
| *Atalopedes campestris* | *Boloria chariclea* | *Callophrys irus* |
| *Atrytonopsis hianna* | *Boloria epithore* | *Callophrys johnsoni* |
| †*Atrytonopsis ovinia* | *Boloria eunomia* | **Callophrys lanoraieensis* |
| *Calpodes ethlius* | *Boloria freija* | **Callophrys mossii* |
| *Carterocephalus palaemon* | *Boloria frigga* | **Callophrys nelsoni* |
| †*Carterocephalus silvicola* | *Boloria improba* | *Callophrys niphon* |
| *Epargyreus clarus* | *Boloria natazhati* | *Callophrys polios* |
| *Erynnis afranius* | *Boloria polaris* | †*Callophrys rubi* |
| **Erynnis baptisiae* | *Boloria selene* | *Callophrys sheridanii* |
| *Erynnis brizo* | †*Cercyonis meadii* | *Callophrys spinetorum* |
| *Erynnis funeralis* | *Cercyonis oetus* | *Calycopis cecrops* |
| *Erynnis horatius* | *Cercyonis pegala* | *Celastrina echo* |
| *Erynnis icelus* | *Cercyonis sthenele* | *Celastrina ladon* |
| *Erynnis juvenalis* | *Chlosyne acastus* | *Celastrina lucia* |
| *Erynnis lucilius* | *Chlosyne damoetas* | *Celastrina neglecta* |
| *Erynnis martialis* | *Chlosyne gorgone* | *Celastrina serotina* |
| †*Erynnis montanus* | *Chlosyne harrisii* | *Cupido amyntula* |
| *Erynnis pacuvius* | *Chlosyne hoffmanni* | *Cupido comyntas* |

*Erynnis persius*
*Erynnis propertius*
†*Erynnis tristis*
*Erynnis zarucco
*Euphyes bimacula
*Euphyes conspicua
*Euphyes dion
*Euphyes dukesi
†*Euphyes peneia*
*Euphyes vestris*
*Hesperia assiniboia*
*Hesperia colorado
*Hesperia comma*
*Hesperia dacotae*
†*Hesperia florinda*
*Hesperia juba*
*Hesperia leonardus*
*Hesperia nevada*
*Hesperia ottoe
*Hesperia pahaska
*Hesperia sassacus*
*Hesperia uncas
*Hylephila phyleus*
*Lerema accius*
*Megathymus streckeri*
*Notamblyscirtes simius*
*Oarisma garita*
*Oarisma powesheik
†*Ochlodes ochracea*
*Ochlodes sylvanoides*
*Panoquina ocola*
*Chlosyne nycteis*
*Chlosyne palla*
*Chlosyne whitneyi*
*Coenonympha nipisiquit
*Coenonympha tullia*
*Danaus plexippus*
†*Erebia cassioides*
*Erebia disa*
*Erebia discoidalis*
*Erebia epipsodea*
†*Erebia euryale*
*Erebia fasciata*
*Erebia lafontainei*
†*Erebia ligea*
*Erebia mackinleyensis*
*Erebia magdalena*
*Erebia mancinus*
*Erebia occulta*
*Erebia pawlowskii*
*Erebia rossii*
*Erebia vidleri*
*Erebia youngi*
*Euphydryas anicia*
*Euphydryas chalcedona*
*Euphydryas editha*
*Euphydryas gillettii*
*Euphydryas phaeton*
*Euptoieta claudia*
*Euptoieta hegesia*
*Junonia coenia*
*Lethe anthedon*
*Echinargus isola*
†*Erora badeta*
*Erora laeta
*Euphilotes ancilla*
*Euphilotes battoides*
*Euphilotes enoptes*
*Feniseca tarquinius*
*Glaucopsyche lygdamus*
*Glaucopsyche piasus*
*Icaricia acmon*
*Icaricia icarioides*
*Icaricia lupini*
*Icaricia saepiolus*
*Icaricia shasta*
*Leptotes marina*
*Lycaena cupreus*
*Lycaena dione*
*Lycaena dorcas*
*Lycaena dospassosi*
*Lycaena editha*
*Lycaena epixanthe*
*Lycaena helloides*
*Lycaena heteronea*
*Lycaena hyllus*
*Lycaena mariposa*
*Lycaena nivalis*
*Lycaena phlaeas*
*Lycaena rubidus*
*Parrhasius m-album
†*Parrhasius polibetes*
*Plebejus anna*

*Pholisora catullus*
*Poanes hobomok*
**Poanes massasoit*
**Poanes viator*
*Poanes zabulon*
**Polites baracoa*
*Polites draco*
*Polites mystic*
*Polites origenes*
*Polites peckius*
*Polites rhesus*
*Polites sabuleti*
**Polites sonora*
*Polites themistocles*
*Polites vibex*
*Pompeius verna*
*Pyrgus centaureae*
*Pyrgus communis*
*Pyrgus ruralis*
*Pyrgus scriptura*
†*Staphylus ceos*
**Staphylus hayhurstii*
**Thorybes bathyllus*
*Thorybes pylades*
*Thymelicus lineola*
*Urbanus proteus*
*Wallengrenia egeremet*

**Lethe appalachia*
†*Lethe diana*
*Lethe eurydice*
†*Lethe minerva*
*Libytheana carinenta*
*Limenitis archippus*
*Limenitis arthemis*
*Limenitis lorquini*
*Limenitis weidemeyerii*
*Megisto cymela*
*Neominois ridingsii*
*Nymphalis antiopa*
*Nymphalis californica*
*Nymphalis l-album*
*Oeneis alberta*
*Oeneis alpina*
*Oeneis bore*
*Oeneis chryxus*
*Oeneis jutta*
*Oeneis macounii*
*Oeneis melissa*
*Oeneis nevadensis*
†*Oeneis norna*
*Oeneis philipi*
*Oeneis polixenes*
*Oeneis uhleri*
*Phyciodes batesii*
*Phyciodes cocyta*
*Phyciodes mylitta*
*Phyciodes pallida*
*Phyciodes pulchella*
*Phyciodes tharos*

*Plebejus idas*
*Plebejus melissa*
*Polyommatus icarus*
**Satyrium acadica*
*Satyrium behrii*
*Satyrium calanus*
**Satyrium californica*
**Satyrium caryaevorous*
**Satyrium edwardsii*
*Satyrium favonius*
**Satyrium fuliginosa*
*Satyrium liparops*
†*Satyrium pruni*
**Satyrium saepium*
**Satyrium semiluna*
*Satyrium sylvinus*
*Satyrium titus*
†*Satyrium w-album*
*Strymon melinus*

| | |
|---|---|
| | *Polygonia comma* |
| | *Polygonia faunus* |
| | *Polygonia gracilis* |
| | *Polygonia interrogationis* |
| | *Polygonia oreas* |
| | *Polygonia progne* |
| | *Polygonia satyrus* |
| | *Speyeria aphrodite* |
| | *Speyeria atlantis* |
| | *Speyeria callippe* |
| | **Speyeria coronis* |
| | *Speyeria cybele* |
| | †*Speyeria diana* |
| | *Speyeria edwardsii* |
| | **Speyeria hesperis* |
| | *Speyeria hydaspe* |
| | *Speyeria idalia* |
| | *Speyeria mormonia* |
| | *Speyeria zerene* |
| | *Vanessa annabella* |
| | *Vanessa atalanta* |
| | *Vanessa cardui* |
| | *Vanessa virginiensis* |
| **Papilionidae (Family)** | **Pieridae (Family)** |
| *Battus philenor* | *Abaeis nicippe* |
| *Eurytides marcellus* | †*Anthocharis belia* |
| *Papilio brevicauda* | *Anthocharis sara* |
| *Papilio canadensis* | †*Anthocharis scolymus* |
| *Papilio cresphontes* | **Anthocharis stella* |
| **Papilio eurymedon* | *Ascia monuste* |
| *Papilio glaucus* | *Colias alexandra* |
| *Papilio indra* | *Colias canadensis* |

| | |
|---|---|
| *Papilio machaon* | *Colias christina* |
| *Papilio multicaudata* | *Colias eurytheme* |
| *Papilio polyxenes* | *Colias gigantea* |
| *Papilio rutulus* | *Colias hecla* |
| *Papilio troilus* | *Colias interior* |
| *Papilio zelicaon* | **Colias johanseni* |
| *Parnassius clodius* | *Colias meadii* |
| *Parnassius eversmanni* | *Colias meadii elis* |
| *Parnassius phobeus* | *Colias nastes* |
| *Parnassius smintheus* | *Colias occidentalis* |
| | *Colias palaeno* |
| | *Colias pelidne* |
| | *Colias philodice* |
| | *Colias tyche* |
| | *Euchloe ausonides* |
| | *Euchloe creusa* |
| | †*Euchloe hyantis* |
| | *Euchloe lotta* |
| | *Euchloe naina* |
| | *Euchloe olympia* |
| | *Eurema mexicana* |
| | *Nathalis iole* |
| | *Neophasia menapia* |
| | *Phoebis philea* |
| | *Phoebis sennae* |
| | *Pieris angelika* |
| | *Pieris marginalis* |
| | *Pieris oleracea* |
| | *Pieris rapae* |
| | *Pieris virginiensis* |
| | *Pontia beckerii* |

| | | |
|---|---|---|
| | *Pontia occidentalis*<br>*Pontia protodice*<br>*Pontia sisymbrii*<br>*Pyrisitia lisa*<br>*Zerene cesonia* | |
| ***Amblyscirtes*** **(Genus)** | ***Anthocharis*** **(Genus)** | ***Callophrys*** **(Genus)** |
| †*Amblyscirtes exoteria*<br>**Amblyscirtes hegon*<br>**Amblyscirtes oslari*<br>*Amblyscirtes vialis* | †*Anthocharis belia*<br>*Anthocharis sara*<br>†*Anthocharis scolymus*<br>**Anthocharis stella* | *Callophrys affinis*<br>**Callophrys augustinus*<br>†*Callophrys chalybeitincta*<br>**Callophrys eryphon*<br>**Callophrys fotis*<br>*Callophrys gryneus*<br>*Callophrys henrici*<br>*Callophrys irus*<br>*Callophrys johnsoni*<br>**Callophrys lanoraieensis*<br>**Callophrys mossii*<br>**Callophrys nelsoni*<br>*Callophrys niphon*<br>*Callophrys polios*<br>†*Callophrys rubi*<br>*Callophrys sheridanii*<br>*Callophrys spinetorum* |
| ***Coenonympha*** **(Genus)** | ***Colias*** **(Genus)** | ***Erora*** **(Genus)** |
| **Coenonympha nipisiquit*<br>*Coenonympha tullia* | *Colias alexandra*<br>*Colias canadensis*<br>*Colias christina*<br>*Colias eurytheme*<br>*Colias gigantea*<br>*Colias hecla*<br>*Colias interior* | †*Erora badeta*<br>**Erora laeta* |

| | | |
|---|---|---|
| | **Colias johanseni* | |
| | *Colias meadii* | |
| | *Colias meadii elis* | |
| | *Colias nastes* | |
| | *Colias occidentalis* | |
| | *Colias palaeno* | |
| | *Colias pelidne* | |
| | *Colias philodice* | |
| | *Colias tyche* | |
| ***Erynnis* (Genus)** | ***Euphyes* (Genus)** | ***Hesperia* (Genus)** |
| *Erynnis afranius* | **Euphyes bimacula* | *Hesperia assiniboia* |
| **Erynnis baptisiae* | **Euphyes conspicua* | **Hesperia colorado* |
| *Erynnis brizo* | **Euphyes dion* | *Hesperia comma* |
| *Erynnis funeralis* | **Euphyes dukesi* | *Hesperia dacotae* |
| *Erynnis horatius* | *†Euphyes peneia* | *†Hesperia florinda* |
| *Erynnis icelus* | *Euphyes vestris* | *Hesperia juba* |
| *Erynnis juvenalis* | | *Hesperia leonardus* |
| *Erynnis lucilius* | | *Hesperia nevada* |
| *Erynnis martialis* | | **Hesperia ottoe* |
| *†Erynnis montanus* | | **Hesperia pahaska* |
| *Erynnis pacuvius* | | *Hesperia sassacus* |
| *Erynnis persius* | | **Hesperia uncas* |
| *Erynnis propertius* | | |
| *†Erynnis tristis* | | |
| **Erynnis zarucco* | | |
| ***Lethe* (Genus)** | ***Oarisma* (Genus)** | ***Oeneis* (Genus)** |
| *Lethe anthedon* | *Oarisma garita* | *Oeneis alberta* |
| **Lethe appalachia* | **Oarisma powesheik* | *Oeneis alpina* |
| *†Lethe diana* | | *Oeneis bore* |
| *Lethe eurydice* | | *Oeneis chryxus* |
| *†Lethe minerva* | | *Oeneis jutta* |

| | | |
|---|---|---|
| | | *Oeneis macounii*<br>*Oeneis melissa*<br>*Oeneis nevadensis*<br>†*Oeneis norna*<br>*Oeneis philipi*<br>*Oeneis polixenes*<br>*Oeneis uhleri* |
| ***Papilio*** **(Genus)** | ***Parnassius*** **(Genus)** | ***Parrhasius*** **(Genus)** |
| *Papilio brevicauda*<br>*Papilio canadensis*<br>*Papilio cresphontes*<br>**Papilio eurymedon*<br>*Papilio glaucus*<br>*Papilio indra*<br>*Papilio machaon*<br>*Papilio multicaudata*<br>*Papilio polyxenes*<br>*Papilio rutulus*<br>*Papilio troilus*<br>*Papilio zelicaon* | *Parnassius clodius*<br>*Parnassius eversmanni*<br>*Parnassius phobeus*<br>*Parnassius smintheus* | **Parrhasius m-album*<br>†*Parrhasius polibetes* |
| ***Poanes*** **(Genus)** | ***Polites*** **(Genus)** | ***Satyrium*** **(Genus)** |
| *Poanes hobomok*<br>**Poanes massasoit*<br>**Poanes viator*<br>*Poanes zabulon* | **Polites baracoa*<br>*Polites draco*<br>*Polites mystic*<br>*Polites origenes*<br>*Polites peckius*<br>*Polites rhesus*<br>*Polites sabuleti*<br>**Polites sonora*<br>*Polites themistocles*<br>*Polites vibex* | **Satyrium acadica*<br>*Satyrium behrii*<br>*Satyrium calanus*<br>**Satyrium californica*<br>**Satyrium caryaevorous*<br>**Satyrium edwardsii*<br>*Satyrium favonius*<br>**Satyrium fuliginosa*<br>*Satyrium lipaprops*<br>†*Satyrium pruni* |

| | | |
|---|---|---|
| | | **Satyrium saepium*<br>**Satyrium semiluna*<br>*Satyrium sylvinus*<br>*Satyrium titus*<br>†Satyrium w-album |
| ***Speyeria*** **(Genus)** | ***Staphylus*** **(Genus)** | ***Thorybes*** **(Genus)** |
| *Speyeria aphrodite*<br>*Speyeria atlantis*<br>*Speyeria callippe*<br>**Speyeria coronis*<br>*Speyeria cybele*<br>†*Speyeria diana*<br>*Speyeria edwardsii*<br>**Speyeria hesperis*<br>*Speyeria hydaspe*<br>*Speyeria idalia*<br>*Speyeria mormonia*<br>*Speyeria zerene* | †*Staphylus ceos*<br>**Staphylus hayhurstii* | **Thorybes bathyllus*<br>*Thorybes pylades* |
